# MuscleNET: mapping electromyography to kinematic and dynamic biomechanical variables by machine learning

**DOI:** 10.1101/2021.07.07.451532

**Authors:** Ali Nasr, Sydney Bell, Jiayuan He, Rachel L. Whittaker, Ning Jiang, Clark R. Dickerson, John McPhee

## Abstract

**Objective:** This paper proposes machine learning models for mapping surface electromyography (sEMG) signals to regression of joint angle, joint velocity, joint acceleration, joint torque, and activation torque.

**Approach:** The regression models, collectively known as MuscleNET, take one of four forms: ANN (Forward Artificial Neural Network), RNN (Recurrent Neural Network), CNN (Convolutional Neural Network), and RCNN (Recurrent Convolutional Neural Network). Inspired by conventional biomechanical muscle models, delayed kinematic signals were used along with sEMG signals as the machine learning model’s input; specifically, the CNN and RCNN were modeled with novel configurations for these input conditions. The models’ inputs contain either raw or filtered sEMG signals, which allowed evaluation of the filtering capabilities of the models. The models were trained using human experimental data and evaluated with different individual data.

**Main results:** Results were compared in terms of regression error (using the root-mean-square) and model computation delay. The results indicate that the RNN (with filtered sEMG signals) and RCNN (with raw sEMG signals) models, both with delayed kinematic data, can extract underlying motor control information (such as joint activation torque or joint angle) from sEMG signals in pick-and-place tasks. The CNNs and RCNNs were able to filter raw sEMG signals.

**Significance:** All forms of MuscleNET were found to map sEMG signals within 2 ms, fast enough for real-time applications such as the control of exoskeletons or active prostheses. The RNN model with filtered sEMG and delayed kinematic signals is particularly appropriate for applications in musculoskeletal simulation and biomechatronic device control.

## 1. Introduction

For volitional control, rehabilitation assessment, intent detection, and power assist, biological signals are the primary inputs of robotic prostheses, active exoskeletons, human-computer interfaces, and rehabilitation robots [1]. One of the primary bio-signals for motion-intention recognition is the electromyography (EMG) signal, which measures the electric potential difference triggered by the nervous system to command muscle contraction [2]. Myoelectric control of biomechatronic devices has encouraged the study of more advanced sEMG control algorithms [3]. Accordingly, two main strategies have been used: (a) detailed muscle modeling and (b) machine learning methods.

One way to convert the sEMG signal to the muscle joint torque is to use a biomechanical muscle model. The most commonly used model is the three-component model, which takes inspiration from the lumped-parameter model developed by A.V. Hill for active and passive muscle tension behavior [4]. However, including a general muscle model within a multibody model introduces various shortcomings such as muscle redundancy, a need to specify complex musculoskeletal geometry such as intricate muscle wrapping pathways and, difficult-to-fit parameters for each muscle, along with sensitivity to these additional parameters. One method that alleviates the muscle geometry, complexity, redundancy, and interpretation of sEMG signals within a control framework uses a machine learning model trained by experimental data.

The output of the machine learning method can be categorized into two groups: (A) a set of decisions, classes, or conditions [5–12], and (B) a data-driven kinematic or kinetic prediction value (regression-based) [13, 14]. The set of classes is usually used for state control and requires a large window of data to make the precise prediction class. Specifically, classification is a post-processing technique that requires the motion data after the motion task is completed. Thus, the regression method has superiority over classification method in terms of real-time processing. The regression-based schemes provide the capability of independent, simultaneous, real-time, and volitional control of each degree of freedom (DoF).

From the general input perspective, the machine learning models fall into two groups: (1) the single instantaneous value of the sEMG signal [15] and (2) pattern recognition techniques (discovering regularities in the pattern of the data) [11, 16–18] Due to the sEMG signals’ stochastic and time-varying nature, sEMG signal windows are utilized for pattern recognition instead of a single instantaneous value. The insufficient and stochastic nature of raw sEMG signals may be solved partially by using a considerable amount of training data and a complex machine learning model.

Transformation of a set or a window of sEMG data into a more readily implemented decreased set of features often uses two methods: (1) feature engineering and (2) machine learning. Feature engineering method’s performance relies on the choice of features for extracting discriminative information from the sEMG data [9, 19–24]. Since the sEMG signal contains temporal or spectral information, the learning algorithms should not rely on specific and limited features. On the other hand, the deep-learning solution uses a hierarchy representation, which learns complex features by configuring and extracting stacks of features [25]. Deep learning architectures, like Recurrent Neural Networks (RNN)s, Convolutional Neural Networks (CNN)s, and Recurrent CNNs (RCNN)s, are primarily used in image analysis or processing [26], speech recognition [27], and recently, bioinformatics [28, 29].

Owing to the CNN’s broad feature learning capability, they have become the most popular deep learning architectures that can perform classification or regression using multi-dimensional data [30]. Lately, CNNs have been applied in sEMG-based classification of hand or wrist gestures [5, 6, 12, 17, 31, 32], and limitedly in sEMG-based regression [33, 34]. Bao et al. [33] used a CNN for wrist multi-DoFs kinematics estimation. Ameri et al. [35] introduced a regression-based CNN that was developed for real-time sEMG based estimation of simultaneous wrist motions.

The RCNN is a particular type of CNN and can mine sequential data or time-frequency information. Chen et al. [13] used an RCNN as sEMG-to-Force mapping for multi-DoFs finger force prediction. They made significant improvements to the prediction results with an RCNN consisting of Long Short-Term Memory (LSTM) networks. Moreover, an RCNN has been utilized for offline estimation of upper-limb motions using sEMG frequency bands [36, 37] and wrist motion intention recognition using the time-frequency spectrum of sEMG signals [38].

The main similarity of the previous studies is that the estimators depend only on the sEMG signal as an input. However, by considering the biomechanical muscle model, the muscle tension relies on the joint’s kinematics in addition to the sEMG signals that function as an activation. The joint angle, velocity, and acceleration partially define the muscle’s tension, with the joint angle specifically defining the muscle fiber orientation. The fiber orientation indicates the direction of tension applied on the bone, and consequently the joint torque. Thus, we propose that in addition to the sEMG signal, the joint’s kinematics be used as input signals to a machine learning model.

In summary, this work’s distinguishing characteristics and novelties are (1) regression-based mapping for intent recognition, muscle modeling, and volitional control of biomechatronic devices, such as exoskeletons, prostheses, and assistive/resistive robots; (2) real-time mapping of sEMG signals by using optimum layers in the machine learning model; (3) proposing a novel structure of CNN and RCNN for filtering raw sEMG signals and feature learning of muscle dynamics; and (4) using delayed joint kinematics as a second input type, inspired by biomechanical muscle models. The proposed MuscleNET (defined as the machine learning muscle models of sEMG signals to regression of kinematic and dynamic biomechanical variables) has several applications, including representing muscles in a musculoskeletal simulation, sports biomechanics simulation, controlling active exoskeletons and prostheses, developing model-based assistive/resistive robots, and post-rehabilitation analysis.

In this paper, firstly, the data preparation steps are described. Secondly, the configuration of the machine learning models is introduced. Finally, the training results of 80 models are discussed and compared.

## 2. Data Preparation

This section presents data collection [39, 40] and processing methods. The data preparation steps have been visualized in Figure 1. The processed data was used for training the machine learning models, as detailed in section 3. The data is available upon reasonable request from the corresponding author [39, 40]. Due to ethical and privacy restrictions, the data is not publicly available.

**Figure 1.**
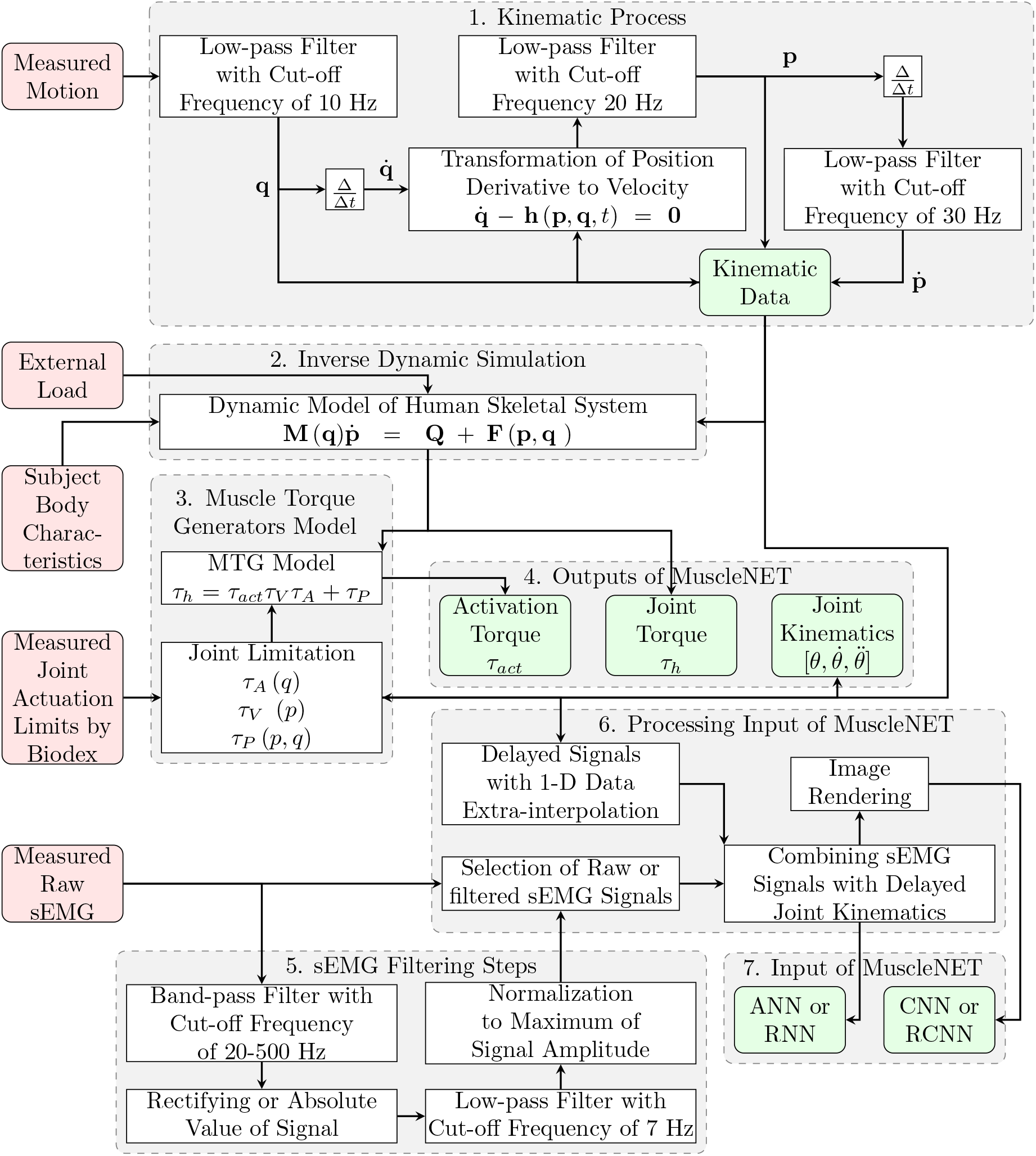
Schematic of data preparation for training of MuscleNET. The inputs of MuscleNET are delayed kinematics and raw and filtered sEMG signals. The output of the MuscleNET may be chosen from the joint angle *θ*, joint velocity 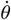, joint acceleration 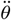, joint torque *τ_h_*, or activation torque *τ_act_*.

### 2.1. Subjects

Seventeen healthy right-handed young individuals (9 Females and 8 Males; 23 ± 4 years; 1.66 ± 0.16 (m) height; 72.25±29.85 (kg) mass) free of upper extremity injury provided informed consent and performed the experimental tasks. The university office of research ethics approved the data collection study.

### 2.2. Instrumentation

Surface EMG signals were measured from 11 sites over muscles of the right upper-limb, similar to those suggested by Avers et al. [41]: the Serratus Anterior (SERR), Middle Deltoid (MDEL), Supraspinatus (SUPR), Infraspinatus (INFR), Posterior Deltoid (PDEL), Pectoralis Major (PECC), Latissimus Dorsi (LATS), Anterior Deltoid (ADEL), Middle Trapezius (MTRA), Upper Trapezius (UTRA), and Lower Trapezius (LTRA). A ground electrode was positioned over the clavicle. Skin sites were shaved and swabbed with an isopropyl alcohol wipe prior to electrode placement. Noraxon Bipolar Surface Ag-AgCl circular electrodes (Noraxon Inc, Arizona, USA) with a fixed 2.0 cm inter-electrode distance were used for placement, and a Noraxon T2000 telemetered system, TeleMyo, (Noraxon Inc, Arizona, USA) was used for collecting signals. We placed the single bipolar electrode over each muscle. Then, we verified signal quality by observing the signal as participants performed isometric contractions in postures that elicit activity in the muscle of interest. Then, the raw sEMG signals were amplified (common-mode rejection ratio ≥ 100 dB at 60 Hz, input impedance 100 MΩ), sampled at a rate of 1500 Hz, and finally, digitalized (16-bit A/D card, maximum ±10 V).

Nine reflective markers and rigid clusters were attached to the subjects on the torso, humerus, and forearm segments. The reflective markers were also installed on targets specific to the task, and the load was controlled by the hand (a bottle). The markers’ 3D position data was recorded by 8 Vicon MX20+ cameras (Vicon Motion Systems, Oxford, U.K.) and sampled at a rate of 50 Hz.

### 2.3. Subject Task Protocol

The data was collected during pick and place tasks [39, 40]. Consequently, the subjects were requested to gradually lift an object from the desk, place it on the upper target, and vice versa, 15 times in 60 seconds (Figure 2). Subjects were asked to rest at the lower and upper target for about 2 seconds. During the mentioned task, the motion of the upper-limb and sEMG signals of muscles were recorded. Participants were asked to sit with their right arm at roughly 30° elevation, 45° external rotation for the thoracohumeral, 120° flexion for the elbow, and the hand grasping the object (the positions of the lower and upper target were adjusted for each participant). The object was a bottle with a 14.9±6.6 (N) weight (each subject used a bottle scaled to their individual shoulder elevation strength, while resulting in different external forces and therefore a more general muscle model [39].)

**Figure 2.**
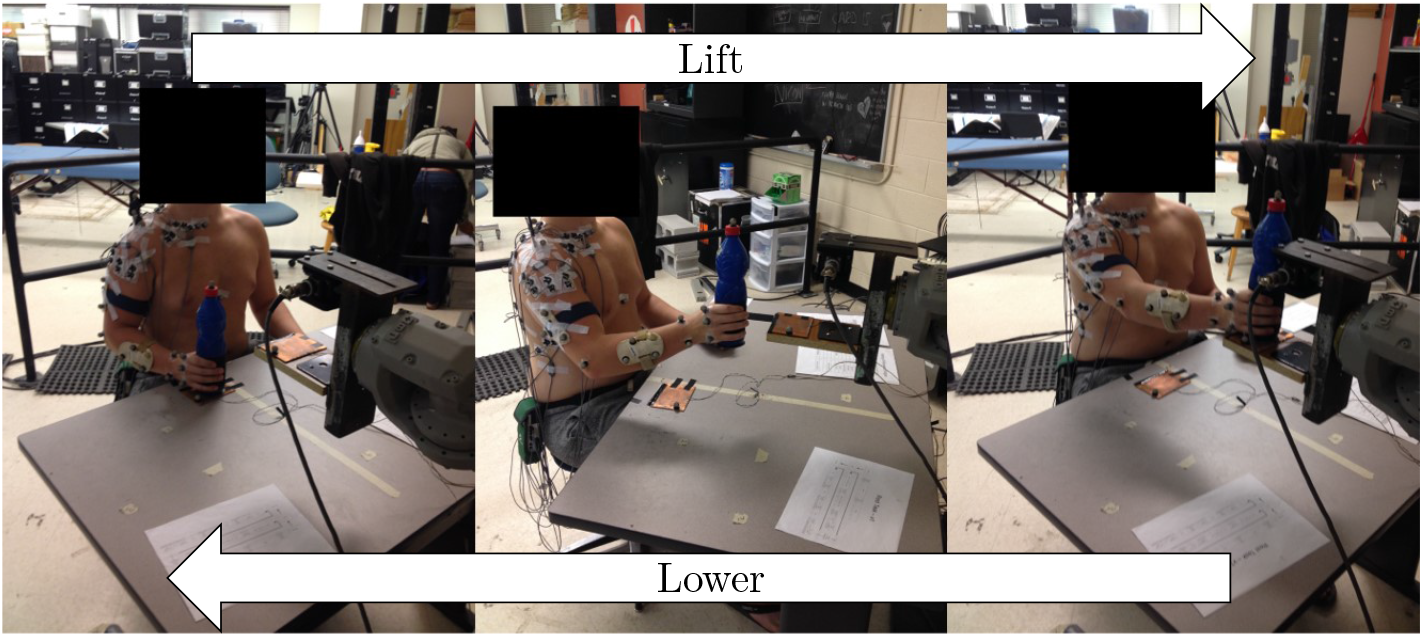
A depiction of the repetitive object manipulation task performed for data collection. The participants lifted and lowered a weighted object between two target locations.

### 2.4. Kinematic Data Analysis and Estimation

Position data was digitally low-pass filtered using a 2^nd^ order Butterworth filter with a cut-off frequency of 10 Hz. Then, re-sampled with the higher sample rate of the sEMG data, 1500 Hz, using 1-D data cubic interpolation of the neighboring grid’s position values with four-position trajectory instances.

The joint angles of the thoracohumeral and forearm were estimated using the cluster markers’ 3D position as well as an anatomical calibration matrix that interprets the correlation between the cluster and coordinate system of the body segment [42]. The angles of the torso to the global and elbow joint were determined according to the International Society of Biomechanics Standard (ISB) recommendations, a Z-X-Y rotation sequence [43]. However, an X-Z-Y sequence (Plane Elevation, Elevation, Axial Rotation) was used for thoracohumeral angles [44] instead of the Y-X-Y recommended by ISB.

The lifting and lowering motions were recognized using the position and velocity of the object. Precisely, the object was estimated at one of the resting targets when its absolute velocity was less than 10 mm/s for more than 40 milliseconds [45].

The joints’ velocity was obtained first using the numerical first derivative of joint angles. Secondly, by the transformation of position derivative to velocity using equation (1). Finally, using a low-pass second-order Butterworth filter with a cut-off frequency of 20 Hz. In the equation (1), **h** is the right-hand side of the transformation between 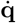 (the vector of all joints position derivatives) and **p** (the vector of all joints velocities).

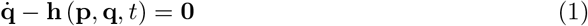

The acceleration of joints was estimated by the numerical first derivative of joint velocities and a low-pass filter using a 2^nd^ order Butterworth filter with a cut-off frequency of 30 Hz.

In this paper, the joint angle, joint velocity, and joint acceleration are defined for the shoulder joint’s elevation.

### 2.5. Delayed Kinematic Data

Since the muscle model relies on the kinematic values as well as the sEMG signal, the kinematic data was used as one of the inputs to MuscleNET. No prior research has combined the kinematic signals with the sEMG signals. Practically acquiring the kinematic values such as angle, velocity, and acceleration has a delay. Thus, we considered a specific amount of delay for these signals. The delay itself has two primary sources: electrical delay and computation delay. The electrical delay relates to the speed of connecting, sampling rate, and computer delay. The computation delay relates to the calculation method delay. For example, since most robots have rotary encoders for measuring the angle, velocity can be calculated using previous angle values with more delay than the joint angle. In other words, the velocity value is not real-time and is based on the previous angle value. In this project, the electrical delay was assumed to be 0.1 s, and the computation delay was 0.05 s, 0.1 s, and 0.15 s for the angle, the velocity, and the acceleration signal, respectively. Thus, in total, the joint angle, joint velocity, and joint acceleration delays of 0.15 s, 0.2 s, and 0.25 s, respectively, from the shoulder joint’s elevation angle may be one set of inputs for the machine learning models. In the biomechatronic control application, the mentioned delay is automatically applied to the signal; thus, it is unnecessary to add additional delay.

The assumed total delay is more than a typical experimental delay because we wanted to analyze the signals in more adverse conditions. The delay for biomechanical simulation is not necessary because real-time performance or processing is not required.

The delayed shoulder elevation joint angle, velocity, and acceleration are *θ* (*t*′), 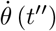 and 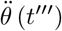, respectively.

### 2.6. Modeling and Inverse Dynamics Simulation

An adapted model performed inverse dynamic simulations. We generated a skeletal model to represent the human upper-limb. We simplified the 3D Stanford VA skeletal arm model [46], which has 10-DoF without the wrist joint. For simplicity, no translational freedom was allowed at the shoulder, only flexion/extension and axial rotation (pronation/supination) at the elbow, and a rigid wrist joint. Thus, our 3D arm model has 5-DoF (three rotations at the shoulder, two rotations at the elbow/forearm). In the 3D model, the shoulder was modeled by three revolute joints with intersecting axes by the Euler coordinate definition. The body segment inertial parameters (BSIP) for the upper-arm (humerus), forearm (ulna and radius), and hand are taken from Dumas et al. [47]. These body segment parameters (dimension, inertia, and joint axes) have been incorporated for modeling by MapleSim. By using the Multibody Analysis Apps of MapleSim, we have extracted motion equations of the skeletal arm as follows:

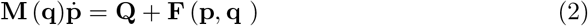

where **M** is the mass matrix, **F** is the right-hand side of the dynamic equations, which consist of Coriolis, centrifugal, and gravitational effects, **Q** is the applied force/torque to the joints, and 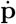 is the vector of all joints’ accelerations. For the given system motion (kinematics), the required forces and torques were calculated by equation (2), which is called inverse dynamic analysis. In this research, the joint torque *τ_h_* is the torque necessary for applying the elevation motion. Specifically, the shoulder elevation joint angle, velocity, and acceleration are *θ*, 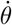, and 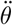, respectively.

### 2.7. MTG (Muscle Torque Generators)

Utilizing machine learning requires significant data (angle /orientation data, velocity data, and external wrench data). Nasr and McPhee showed that using the Muscle Torque Generator (MTG) model [48] requires a smaller amount of data for training the MuscleNET [49]. In other words, these models simulate components of muscle modeling (i.e., orientation constraint of the joint and velocity constraint of the muscle) in joint torque estimation; individual muscles are not explicitly modeled. As an introduction, the MTG model consists of the human joint torque *τ_h_*, the activation torque *τ_act_*, the position-scaling function *τ_a_*, the velocity-scaling function *τ_v_*, and the passive torque function *τ_p_* as shown in equation (3) [48, 50].

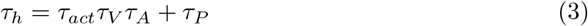

### 2.8. EMG Data Filtering

In this research, the inputs of the MuscleNET are sEMG signals. To evaluate the ability of each model, we have used both raw sEMG and filtered sEMG. The following five steps achieved filtering of the raw signals: 1) A second-order band-pass digital Butterworth filter with a normalized cut-off frequency of 20-500 Hz was used to remove heart rate artifacts and high-frequency content [51, 52] (signals below 20 Hz were cleaned to remove the motion artifacts, and sEMG signals above 500 Hz were cleaned as they had minor power spectral density [53]). 2) A second-order band-stop digital Butterworth filter (notch filtered) with a normalized cut-off frequency of 55-65 Hz was applied to eliminate the 60 Hz noise from the measurement unit. 3) The absolute value of the signal amplitude or rectifying the signal was used. 4) A second-order low-pass digital Butterworth filter with a normalized cut-off frequency of 7 Hz [52] was used to smooth the signal as evaluated and analyzed by Nasr et al. [54]. 5) Normalization to the trial maximum signal amplitude was used. Extra filtering was extraneous since MuscleNET is a machine learning model for mapping signals (not a mathematical muscle model). A sample of the raw and filtered sEMG in the time and frequency domain is shown in Figure 3. Studying signal loss was out of the research scope; therefore, the data measurement was repeated in case of signal disconnection.

**Figure 3.**
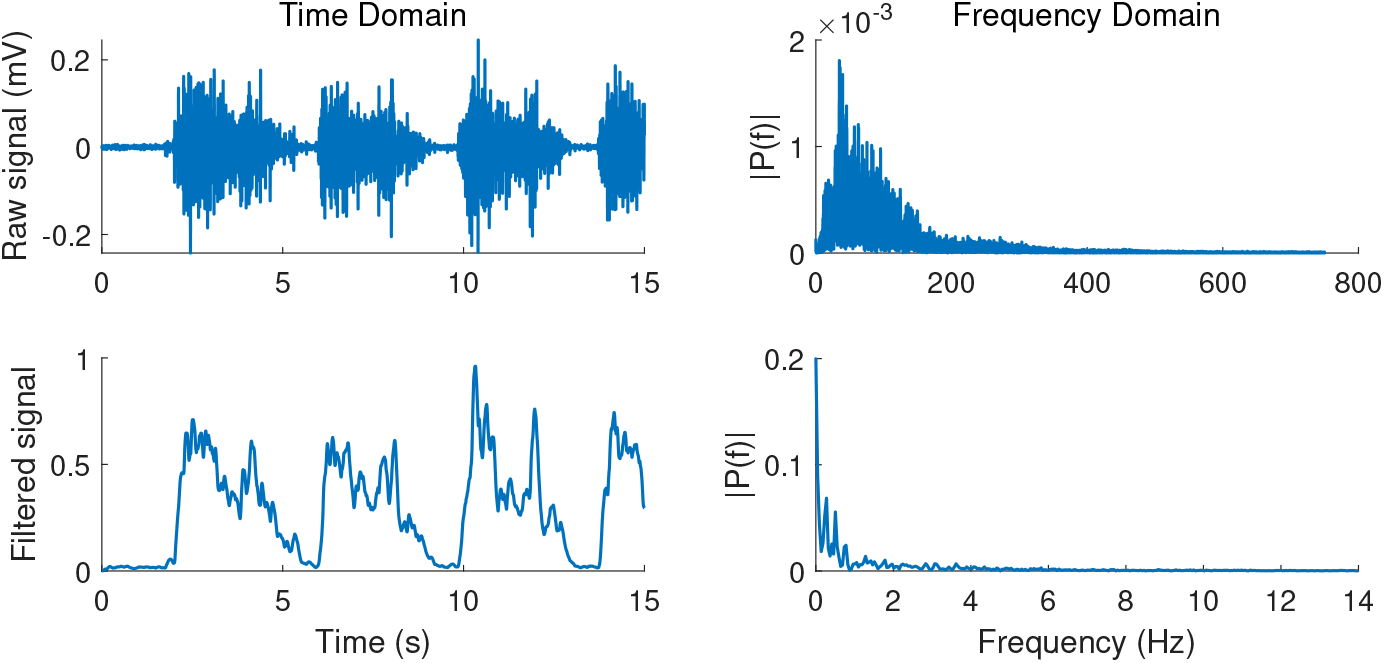
A sample of raw and filtered sEMG signals in the time domain and frequency domain.

## 3. Machine Learning Models

This section describes the four different machine learning models: ANN (Forward Artificial Neural Network), RNN (Recurrent Neural Network), CNN (Convolutional Neural Network), and RCNN (Recurrent Convolutional Neural Network). Configuration differences related to input type are described in detail. The machine learning models’ output can be the joint angle, joint velocity, joint acceleration, joint torque *τ_h_*, and activation torque *τ_act_* signals. The outputs were normalized by the maximum value for each subject. The performance was reported with the computation of:

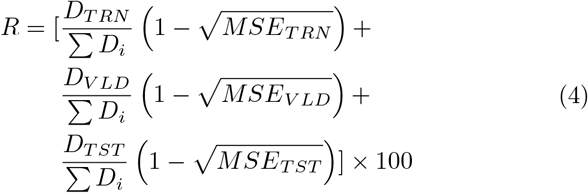

where *MSE* is the mean squared error between the output values and target values, *D* is the amount of data, and the subscripts *TRN*, *VLD*, and *TST* are training, validation, and testing volume, respectively.

The initial architecture of machine learning models was adopted from prior researches [13, 28, 33, 35–37, 55]. The number of hidden layers, neurons in each hidden layer, feedback delay, input delay, convolution filtering size, pooling size, and strides were selected by trial and error (Figure 4) to have the highest regression accuracy. The CNN and RCNN MuscleNET with delayed kinematics have a novel configuration, which are introduced in section 3.3.2.

**Figure 4.**
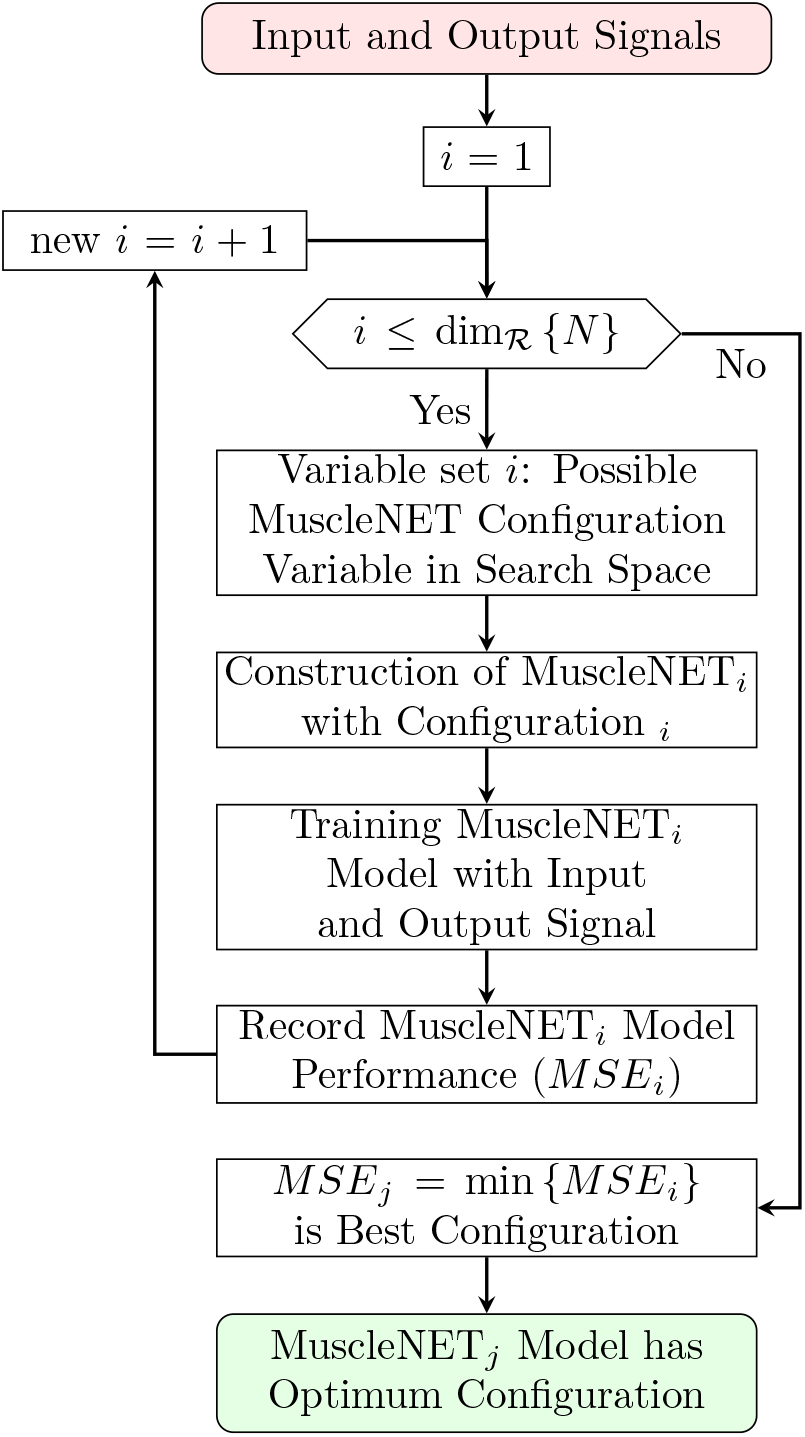
Schematic of MuscleNET models configuration optimization loop.

1560000 pairs of inputs and outputs (1560000 = 17 subjects ×60 seconds ×1500 Hz sample rate) were used for training, validating, and testing the MuscleNET. To evaluate the amount of experimental data required for the machine learning model, a complete estimation was done on the test subject who did not participate in the training and validation of the model. In situations where the model had more than 85% regression accuracy, the capability of tuning the general model to estimate a test subject’s biomechanical variables was demonstrated.

The regression proficiency of the machine learning model depends on the training algorithm (Figure 5). Dividing data for training, validation, and testing is controversial and depends on the nature of the datasets, the number of variables, and the machine learning model architecture. There is no strict rule for dividing, and therefore each method should be evaluated separately as intrinsic characteristics vary (model architecture, data nature, and model purpose). Using more data sets for training and less data for validation and testing can increase the risk of overfitting [56]. We have added a discussion of this issue to the paper in section 4.

**Figure 5.**
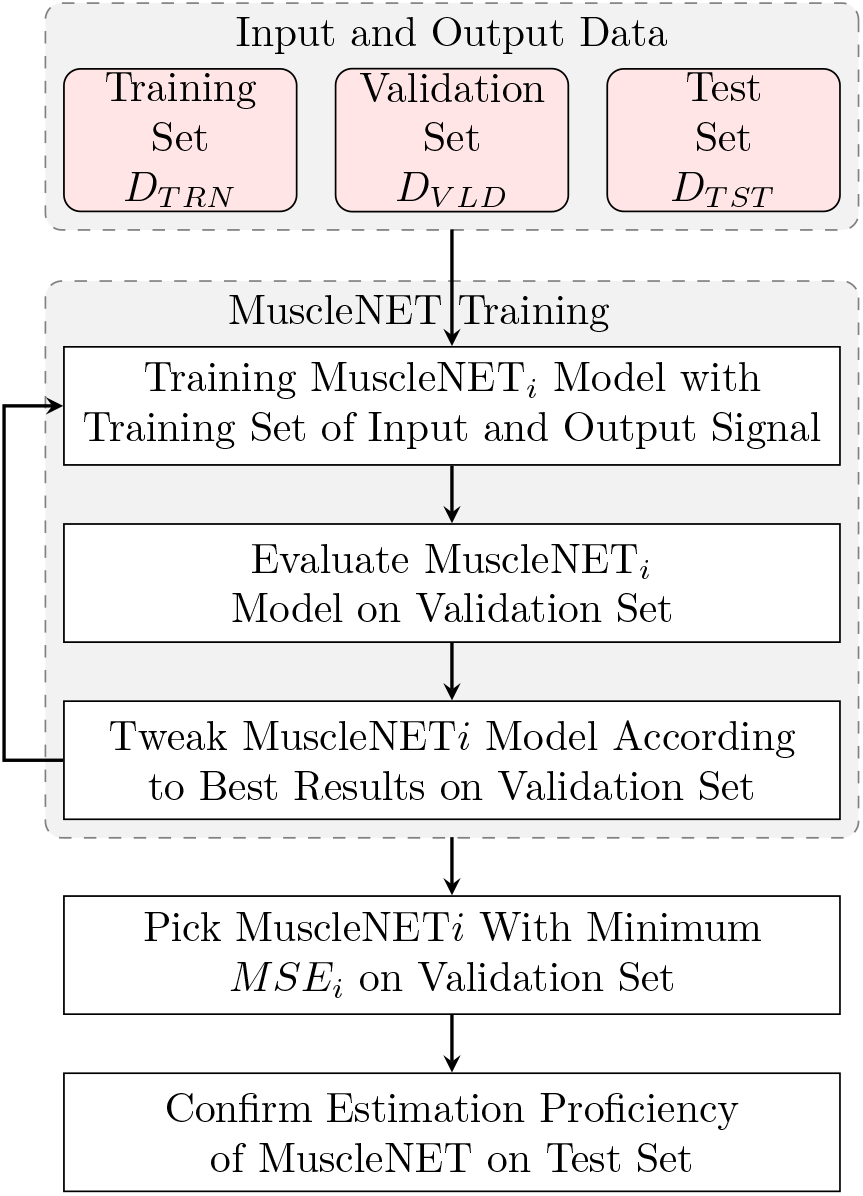
Schematic of training MuscleNET with training, validation, and testing sets of the input and output signal.

### 3.1. ANN (Forward MuscleNET)

One of the first and simplest machine learning models for modeling a muscle is the forward artificial learning model. According to Nasr et al. [55], this multilayer model should be wide and shallow to map the sEMG to activation torque with minimal error and high accuracy. Their recommended architecture [55] was adopted, and different configurations (number of hidden layers and neurons in each hidden layer) were tested using the data sets to obtain the optimized configuration (for maximum accuracy cost function). The optimized configuration had the highest regression accuracy in the same training condition and is described in the following.

If the sEMG signal was raw, the number of hidden layers was set at 2, and if the sEMG was filtered, it was set at 1. One additional layer was added to the Nasr et al. [55] model for the described approach as more layers filtered the raw signal. Since the delayed kinematics provided valuable data for the model, the number of neurons in each layer was set to 4 times the number of input signals (for example, for 11 sEMG signals and 3 delayed kinematic signals, the number of neurons is 4 times 14 or 56). When the inputs were purely sEMG signals, the number of neurons in each layer was set at 5 times the number of input signals (for example, for 11 sEMG signals, the number of neurons is 5 times 11 or 55). ANN MuscleNET configuration details are summarized in Table 1.

**Table 1.** Detail of ANN MuscleNET Model Configuration.

| # | Inputs |  | Configuration |  |  |  | Outputs |  |
| --- | --- | --- | --- | --- | --- | --- | --- | --- |
|  | Filtered sEMG | Size | Number of |  | Activation Function in |  | Size | Number of Neurons |
|  | Raw sEMG |  | Hidden Layers | Neurons in Each Hidden Layer | Hidden Layer | Output Layer |  |  |
| 1 | X | 11×1 | 1 | 55 | Tan-sigmoid | Tan-sigmoid | 1×1 | 716 |
| 2 | X | 11×1 | 2 | 55 | Tan-sigmoid | Tan-sigmoid | 1×1 | 3796 |
| 3 | X X | 14×1 | 1 | 56 | Tan-sigmoid | Tan-sigmoid | 1×1 | 897 |
| 4 | XX | 14×1 | 2 | 56 | Tan-sigmoid | Tan-sigmoid | 1×1 | 4089 |

The training method was a Levenberg-Marquardt backpropagation. The maximum epoch was set to 2000. The maximum elapsed time was set to 6 hours.

### 3.2. RNN (Recurrent MuscleNET)

Since the muscle dynamics and joint motion rely on motion history, we hypothesized that Recurrent Neural Networks might have better accuracy in muscle modeling. This model has feedback from the output to the inputs. After training the nonlinear autoregressive with external input (NARX) networks, the output time series was predicted with the past output values (the feedback input) and the external input time series (sEMG signal and delayed kinematics). The general architecture [13, 28] was adopted, and different configurations (number of hidden layers, neurons in each hidden layer, feedback delays, and input delays) were tested using the data sets to obtain the optimized configuration. The following optimized configuration had the highest regression accuracy in the same training condition.

The number of hidden layers was 2. The number of neurons was 3 times the number of input signals when the inputs had delayed kinematic signals and 4 times the input signals when the inputs did not consist of the delayed kinematics. The feedback delays were then set to the last 7 signal values. Since different input delays did not impact accuracy, we set the input delays to zero or used the current input value. The details of the RNN MuscleNET configuration are summarized in Table 2, and an example of RNN MuscleNET for 11 filtered sEMG signals and 3 delayed kinematic signals is depicted in Figure 6. The maximum epoch training method and the maximum elapsed time were the same as for the ANN.

**Table 2.**
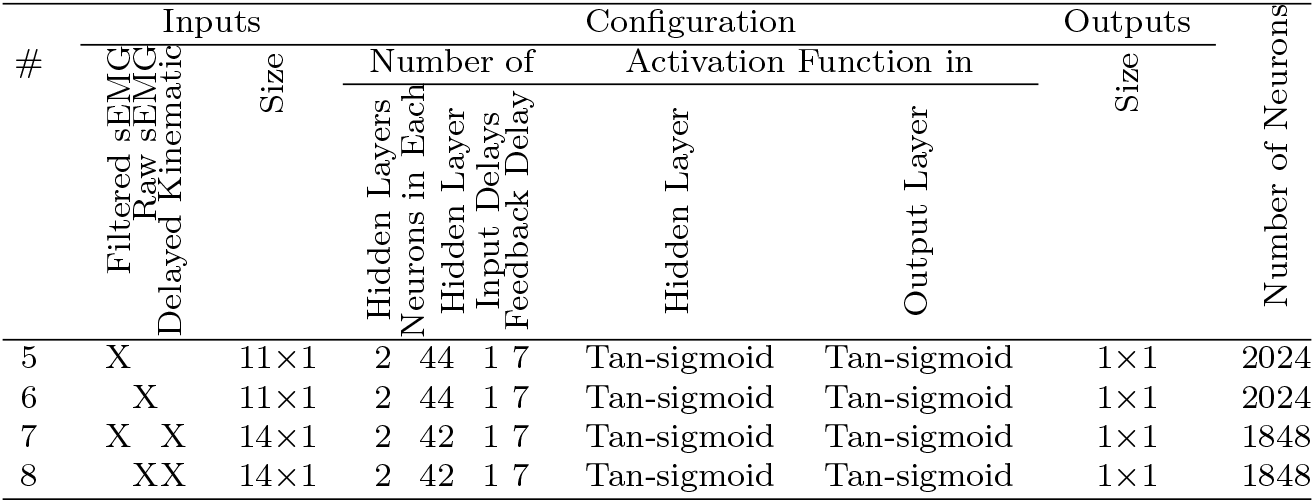
Detail of RNN MuscleNET Model Configuration.

**Figure 6.**
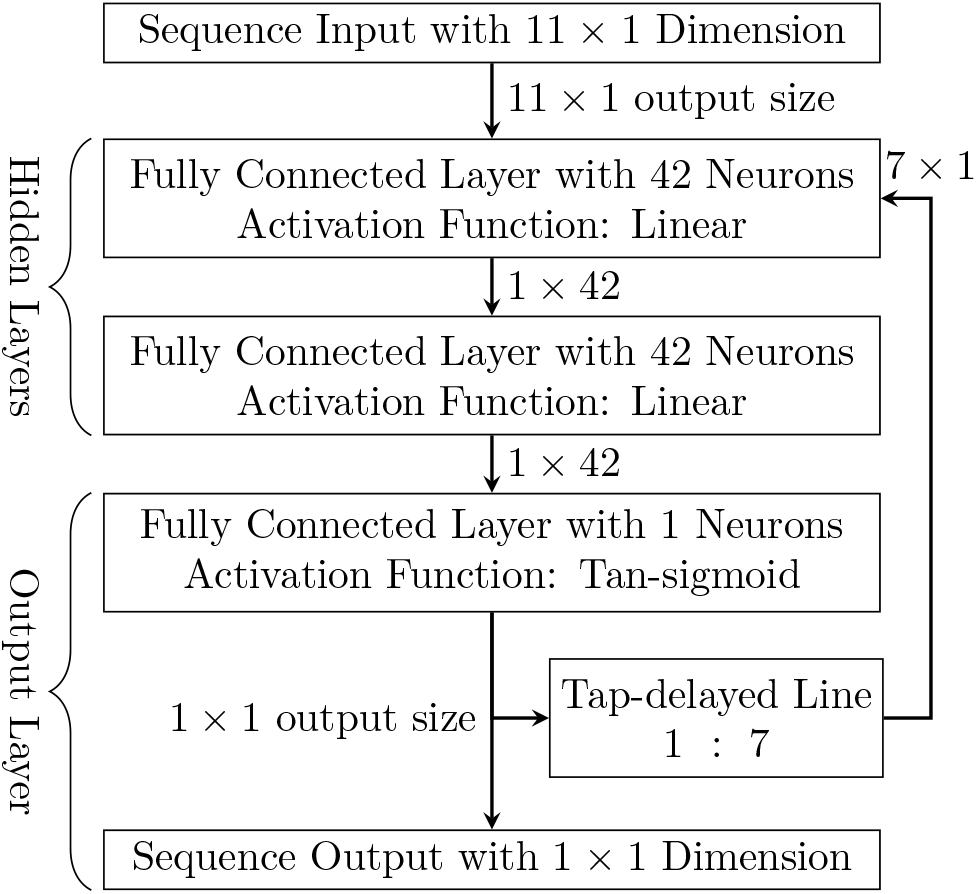
Schematic of the RNN MuscleNET for combined input of 11 filtered sEMG and three delayed kinematic signals.

### 3.3. CNN (Convolutional MuscleNET)

We initially adopted the CNN and RCNN configuration from prior work [31, 33, 37] and tested different configurations (number of convolutional layers, convolution filtering size, pooling size, strides, neurons in fully connected layer, activation function) using the data sets to obtain the following configuration. This configuration had the highest regression accuracy in the same training condition. The configuration of CNN and RCNN MuscleNET with a delayed kinematic signal is novel in terms of parallel structure for delayed kinematic signal input.

Since there were two different input signals (sEMG signals and delayed kinematic signals), the CNN models’ configurations were categorized into 4 different shapes based on the input’s conditions. First, we describe the CNN generic configuration and then the different configurations in the following paragraph.

Generally, the typical configuration consists of an input group, the convolution groups, and an output group. First, the input group is a sequence input layer that gets an image from the input signals. Secondly, the first layer of convolution groups is a folding sequence layer, and the last one is an unfolding sequence layer. Finally, the output group consists of 4 layers: the flatten layer, a fully connected layer with an output size of 50 neurons, a fully connected layer with an output size of 1 neuron, and finally, the regression layer.

Precisely, the number of signals made the image’s height (for example, for 11 sEMG signals and 3 delayed kinematic signals, the image’s height was 14). The image’s width is relevant to the history of sEMG signals that we want to filter by CNN or RCNN. Since the maximum human motion frequency is roughly 6 Hz [57], we recommend using a maximum of 7 Hz. We did not propose to use less than 5 Hz since the number of data points increases for the CNN or RCNN, slowing model processing. For data with a 1500 Hz sample rate, we used 250 points for the image’s width, which is 6 Hz of the data. In summary, the input image size was 11 by 250 for the sEMG signals input and was 14 by 250 for inputs consisting of sEMG signals plus delayed kinematics (Figure 7). The current data is located on the image’s right, and the 249 previous data is located at the left of the image. The sequential input layer handled importing this 1D image. It is noteworthy that signals should not have the same rate as the sampling rate when making the images from the signals. Since the model’s delay might be around 0.05 sec or 20 Hz, we used a slower rate of making images. We used 0.1 sec for making the images, and at each time, must use the current signals and 249 of the previous signals to make the image, then wait for 0.1 sec to make the following image. This way, we considered the model’s delay and system to use the model for model-based control of biomechatronic devices. Using an image composed of sEMG signals and delayed kinematic signals is a novel approach.

**Figure 7.**
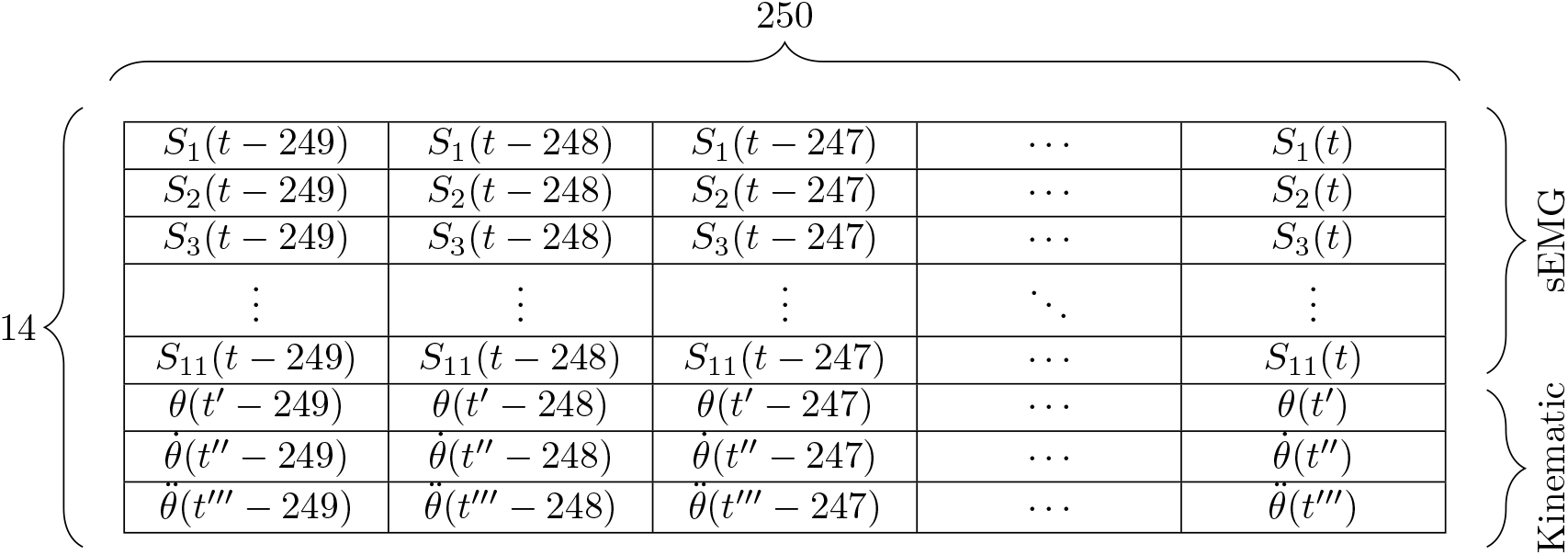
A sample of the input image made from sEMG signals and delayed kinematic signals used as the input of the CNN or RCNN configuration of MuscleNET. The delayed shoulder elevation joint angle, velocity, and acceleration are *θ*(*t*′), 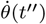, and 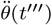, respectively (*t*′, *t*″, and *t*′′′ are the delayed times).

The training method was Adam (adaptive moment estimation) optimizer with a unit threshold for the gradient. The learning rate used for training was set to 0.001. The maximum epoch was set to 500. The mini-batch size (a subset of the training set) for each training iteration was set to 2000.

The convolution groups were unique for the different input signals (with or without delayed kinematics and raw or filtered sEMG signals). In the following sections, the CNN models are explained in detail for each input signal combination.

#### 3.3.1. Without Delayed Kinematic Signals

When only the raw sEMG signals were the input image’s constructor, there were 8 layers in the convolutional groups, and they were taken serially (Figure 8). All groups had a 2-D convolutional layer that was applied to slide convolutional filters with a filter size of height 1 and width 3. The output of this layer had the same size as the input when the stride equaled 1. The step size for traversing the input was 1 vertically and was 1 horizontally. Groups 1, 2, 7, and 8 had 16 neurons, and groups 3 to 6 had 64 neurons in the convolutional layer (the number of channels or feature maps). All 8 groups had a batch normalization layer that normalized each input channel across a mini-batch size. We used this batch normalization layer to accelerate the networks’ learning and reduce the network initialization sensitivity. Moreover, all 8 groups had a Rectified Linear Unit (ReLU) layer that implemented a threshold operation to each input element, where any negative value was set to zero. Only the first 5 groups had average pooling layers for down-sampling by dividing the input into rectangular pooling areas and calculating each area’s average values. The pooling region’s dimensions had a height of 1 and a width of 3. The vertical step size was 1, and the horizontal step size was 3. The vertical unit size for the pooling layer’s stride was utilized because the sEMG signals should be filtered separately.

**Figure 8.**
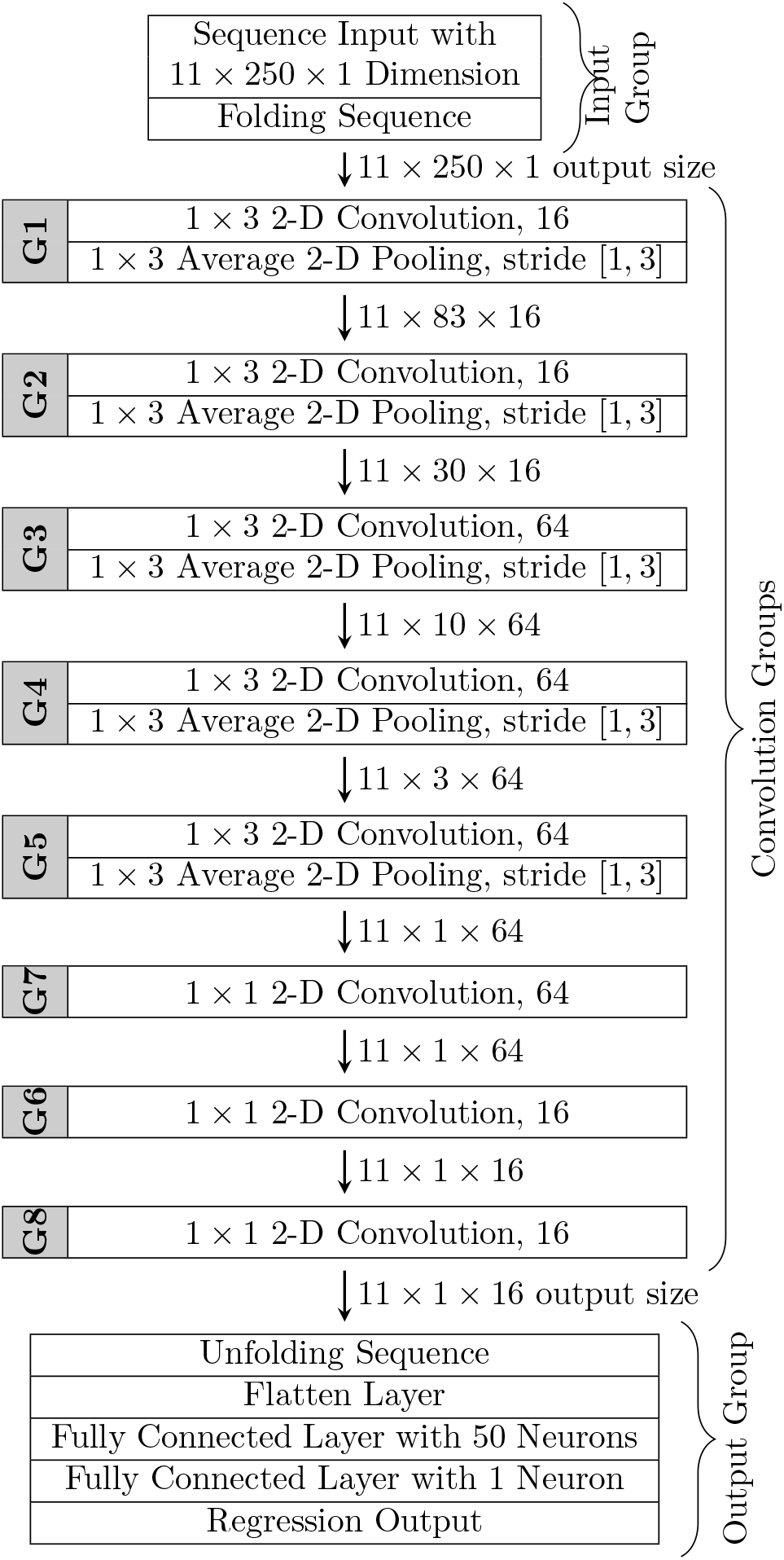
Schematic of the convolutional MuscleNET for raw sEMG signal input only, including convolutional layers, average pooling layers, and fully connected layer. The batch normalization and the rectified linear unit after the convolutional layers in each group have not been shown for simplicity. The convolutional layers and the average pooling layers have the same padding.

All in all, the first 5 groups had a 2-D convolution layer, a batch normalization layer, a ReLU layer, and a 2-D average pooling layer. The 6 to 8 groups had a 2-D convolution layer, a batch normalization layer, and a ReLU layer.

For input images constructed with the filtered sEMG signals, group 7 was removed, and group 6 was connected to group 8. The convolutional neural network configuration for raw sEMG signal and without the delayed kinematics signal was quite similar to the model proposed by Ameri et al. [35], which allowed for a direct comparison with our new model.

#### 3.3.2. With Delayed Kinematic Signals

Since the origin of the delayed kinematic and sEMG signals are different, they first should be filtered with different convolutional layers separately. To this end, we innovated a parallel structure with two separate bridges. The first bridge handles filtering the sEMG signals, and the second one relates to delayed kinematic usage. Finally, both bridges intersect, and all data was used for modeling the mapping of the input (sEMG and delayed kinematic signals) to the output signals. This novel configuration for separating different signals from the image is shown in Figure 9.

**Figure 9.**
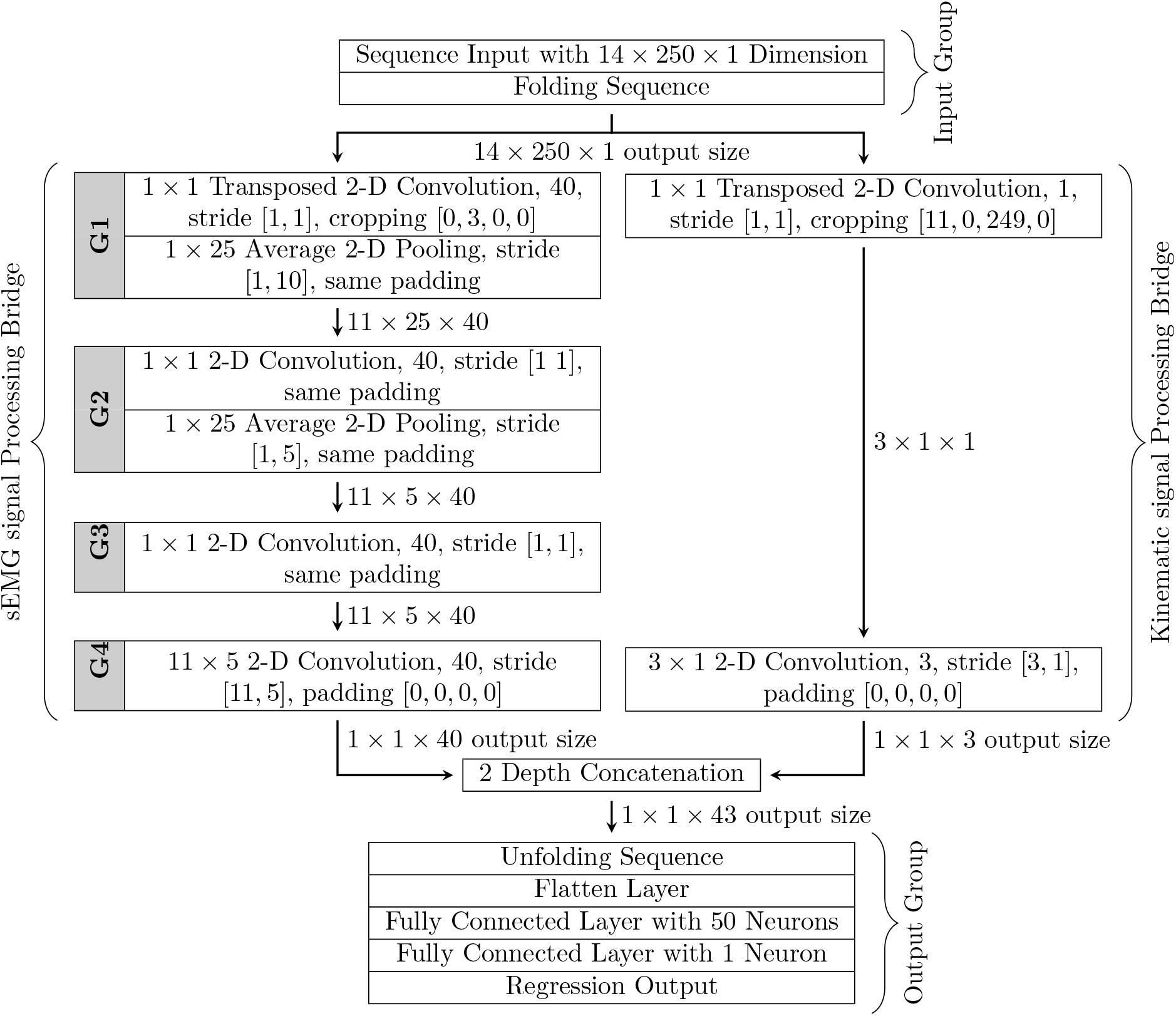
Schematic of the convolutional MuscleNET for combined input of raw sEMG and delayed kinematic signals, including convolutional layers, average pooling layers, and fully connected layer. The batch normalization and the rectified linear unit after the convolutional layers in groups 2 and 3 have not been shown for simplicity.

The left bridge, dedicated to filtering the sEMG signals, had 4 groups for raw sEMG signals. The first group had a transposed 2-D convolution layer with an additional feature of cropping the image. This new feature shaved the input image (that is, the combination of sEMG and delayed kinematic signals). The transposed 2-D convolution layer’s output size was reduced by cropping from the bottom of the input by 3 units since the last 3 pixels are relevant to the delayed kinematic signals. The transposed 2-D convolution layer and 2-D convolutional layers at groups 2 and 3 had a filter size of the height of 1 and the width of 1. All convolutional layers had 40 neurons each. The step size for traversing the input was 1 vertically and was 1 horizontally for all convolutional layers. The first two groups had a ReLU layer and average pooling layers for down-sampling. Both pooling layers had a pooling region’s dimensions with a height of 1 and a width of 25. However, the first one’s stride at the first layer was 10, and the second one at the second layer was 5 in the horizontal direction. The vertical unit size for stride of both pooling layers arose because the sEMG signals should be filtered separately. The last group only had a 2-D convolutional layer with different filter sizes of the height of 11 (since there are 11 sEMG signals) and the width of 5 with the same stride. As shown in Figure 9, since the output size of group 3 was 11×5×40, it is evident that the 250 pixels were converted to 5 pixels. We have tried to have these 5 pixels for pattern detection purposes. Since the sEMG signal was not converted to one signal and had 5 pixels, it can provide more timing information.

For input images constructed with the filtered sEMG signals, group 3 was removed, and group 2 connected to group 4.

The right bridge, which is dedicated to acquiring the delayed kinematic signals, had 2 layers. The first layer had a transposed 2-D convolution layer with a cropping feature from the top of the input image by 11 units (since the first 11 pixels are relevant to the sEMG signals) and from the left of the input image by 249 units (since the last column is the recent signals). Precisely, the cropping converted the 14 by 250 image size to a 3 by 1 image size. The transposed 2-D convolution layer had a filter size of the height of 1 and the width of 1 with 1 neuron. The second layer had a regular 2-D convolutional layer with a filter size of the height of 3 and the width of 1.

The signals from two bridges intersected with a depth concatenation layer. The depth concatenation layer received inputs with the same height of 1 and width of 1 and concatenated them alongside the depth dimension. In this way, both signals of sEMG and delayed kinematics were stacked to an image with 1 width, 1 height, and 43 depth.

### 3.4. RCNN (Recurrent Convolutional MuscleNET)

The RCNN’s configuration approximates the CNN; the only difference is in the output group’s layers. The fully connected layer with an output size of 50 neurons in the CNN was changed to a Long Short-Term Memory (LSTM) layer with 50 hidden units. Additionally, the output of the LSTM was the last time step of the sequence in the RCNN. The activation function to update the cell and hidden state was set to be the hyperbolic tangent function, and the activation function to apply to the gates was the sigmoid.

The schematic of the RCNN MuscleNET for the combined input of raw sEMG and delayed kinematic signals has been shown in Figure 10. The input image, the input group, and the general output group were introduced in section 3.3. The details of the sEMG signal processing and kinematic signal processing have been presented in section 3.3.2.

**Figure 10.**
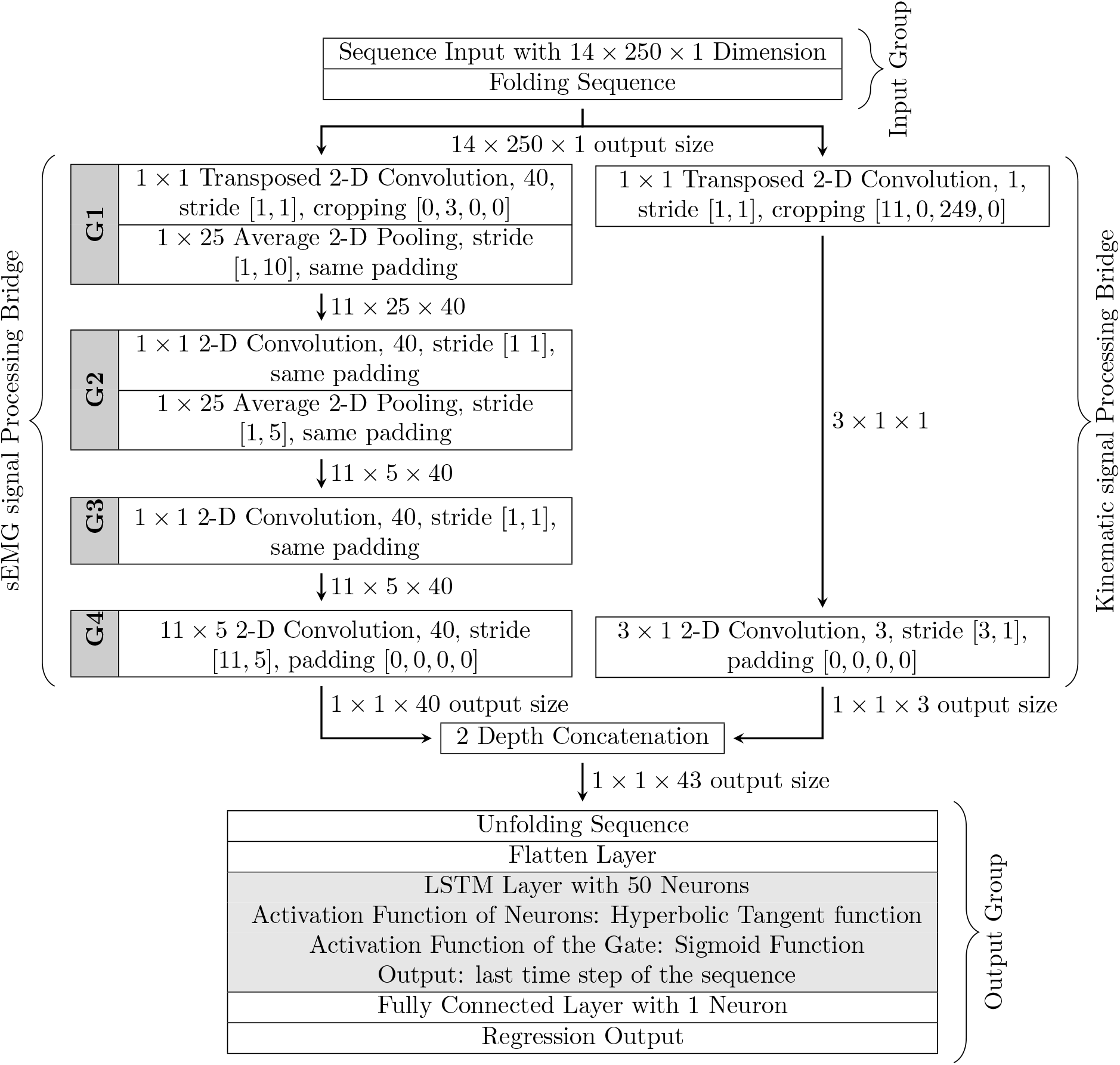
Schematic of the RCNN MuscleNET for combined input of raw sEMG and delayed kinematic signals, including convolutional layers, average pooling layers, and a Long Short-Term Memory (LSTM) layer. The batch normalization and the rectified linear unit after the convolutional layers in groups 2 and 3 have not been shown for simplicity.

## 4. Results and Discussion

This section presents the data ratio analysis for the training, validation, and testing of subject-specific and general models. Second, the volume of experimental data for complete estimation with a general model was assessed. Third, 80 different machine learning configurations of MuscleNET with different input conditions (raw or filtered sEMG signal, without or with delayed kinematic signals) and different outputs (5 different biomechanical signals) were compared in terms of the performance, training time, and maximum inference time. Finally, samples of complete estimation for random subjects were presented. For training of the MuscleNET, a personal computer with an Intel^®^ Core™ i7-3370 CPU @ 3.40GHz processor and 16.0 GB memory was used.

### 4.1. Dividing Ratio and Data Volume

To study the division of data for training, validation, and testing concerning maximum estimation accuracy without overfitting, we trained the models with different ratios. We have assessed the different ratios, and two combinations were ideal for the subject-specific and general models (Figure 11). The subject-specific model was tuned for maximum estimation accuracy of validation data with 94.1% of data for training, 5.9% of data for exporting-and-validation (validation has overlap with training set), and 5.9% unique data for testing. The general model was achieved with 94.1%, 23.5%, and 5.9% of data for training, exporting-and-validation, and testing, respectively (again, the validation sets have overlap with training sets). The overlap of validation and training sets may decrease in the case of reach experimental data availability.

**Figure 11.**
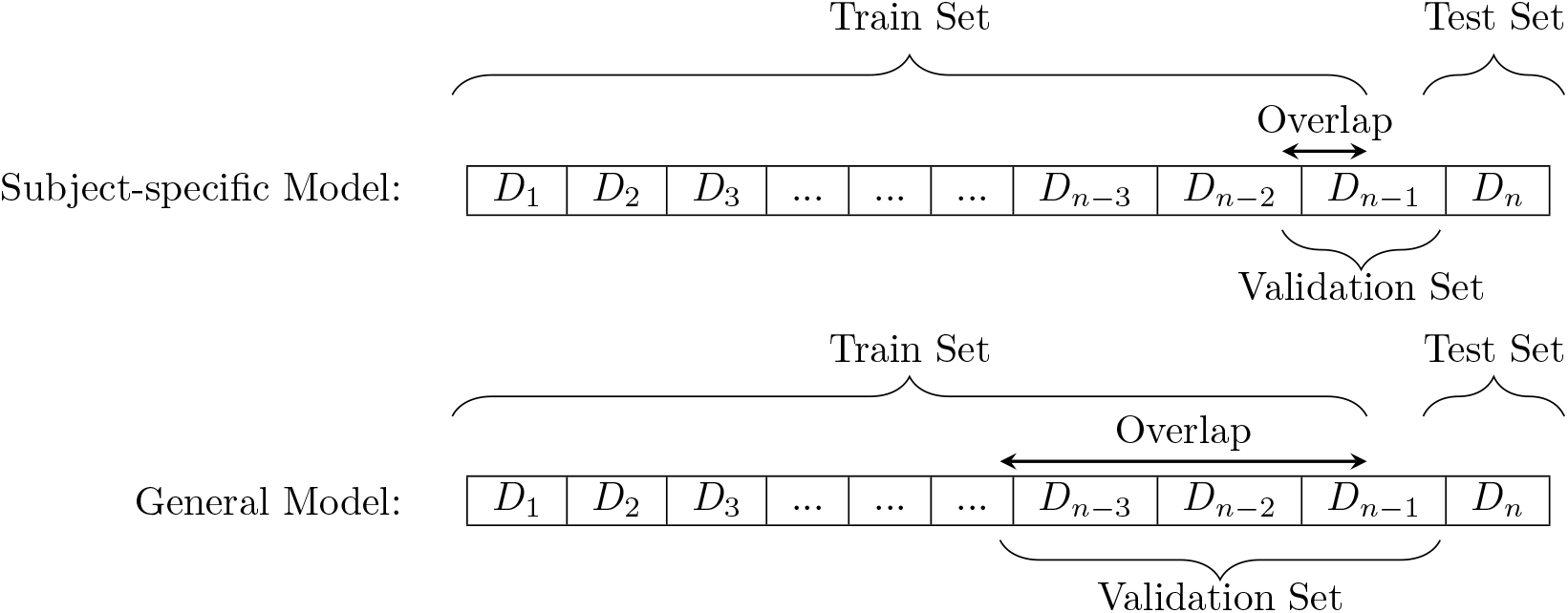
Visual representation of dividing data base into training, validation, and testing sets for subject-specific and general models.

For the subject-specific model, 16 subjects (94.1% of total data) were selected for training, and one of them (5.9% of total) was present in training and validation simultaneously. This subject plays the role of stopping the training process and signaling the export of the final model with the lowest MSE. The other subject which did not participate in training and validation was selected as the test set. We observed that the regression accuracy for the validation subject increased to 99.7% (Figure 12). However, the test result was not entirely successful (97.7%). In other words, since the training stopped and was exported based on the regression of one subject validation, the model is subject-specific for the validation subject and a less general model for testing data sets. The machine learning model has less regression accuracy for testing with different data sets (any subjects other than those used for validation). Although this trained MuscleNET with one validation subject’s data has high regression accuracy for the validation subject, the model is subject-specific and should not be used for general purposes. If the EMG signals were not stochastic and time-varying, the regression would be far more than the amount reached already. In addition, from a biomechanical perspective, the maximum muscle force, muscle attachment configuration, muscle passive and active functions (biomechanical muscle features) are different between different subjects. Thus, selecting a small amount of data for validation (for example, only one subject) decreases the reach and broad biomechanical muscle features of different subjects and increases the validation accuracy for the subject-specific model.

**Figure 12.**
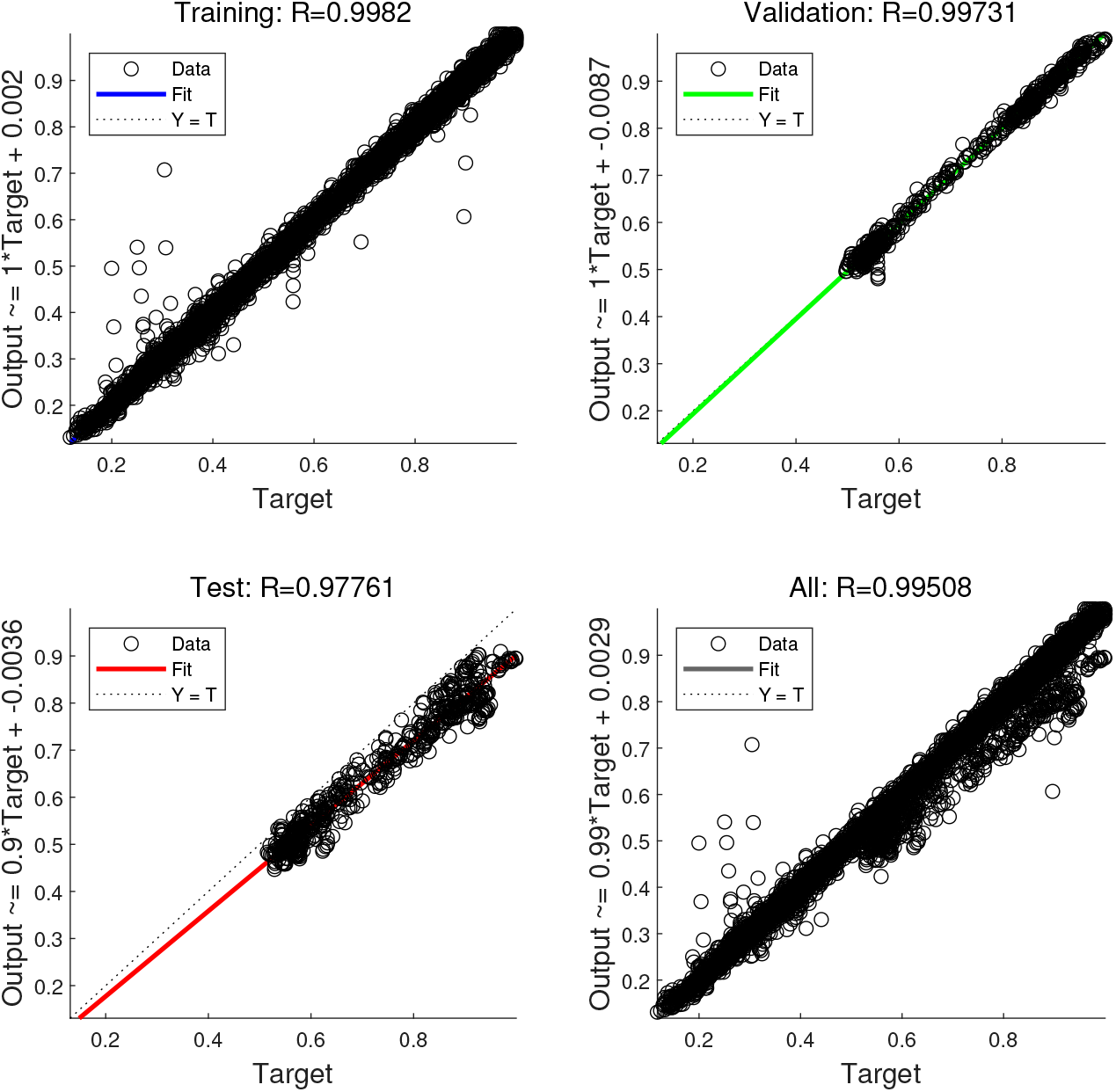
The machine learning training, validation, and testing regression of a subject-specific model of RNN MuscleNET with filtered sEMG and delayed kinematic signals input and joint angle output. The test subject did not participate in training and validation.

For the general model, we found that exporting required 23.5% of data (4 subjects) for validation instead of 5.9% of data (1 subject) for subject-specific cases. The general model estimated the biomechanical signals of the test data set with regression of 99.1% (Figure 13). In other words, less data for validation and more data for training decreased the regression of the general model for testing. This issue is called overfitting in supervised machine learning and prevents generalized machine learning models from properly fitting observed data on training data, along with unobserved data on validation and testing sets [56]. Thus, we proposed that the validation data sets should be more than one person data to have a general machine learning model, ignoring the overfitting issue.

**Figure 13.**
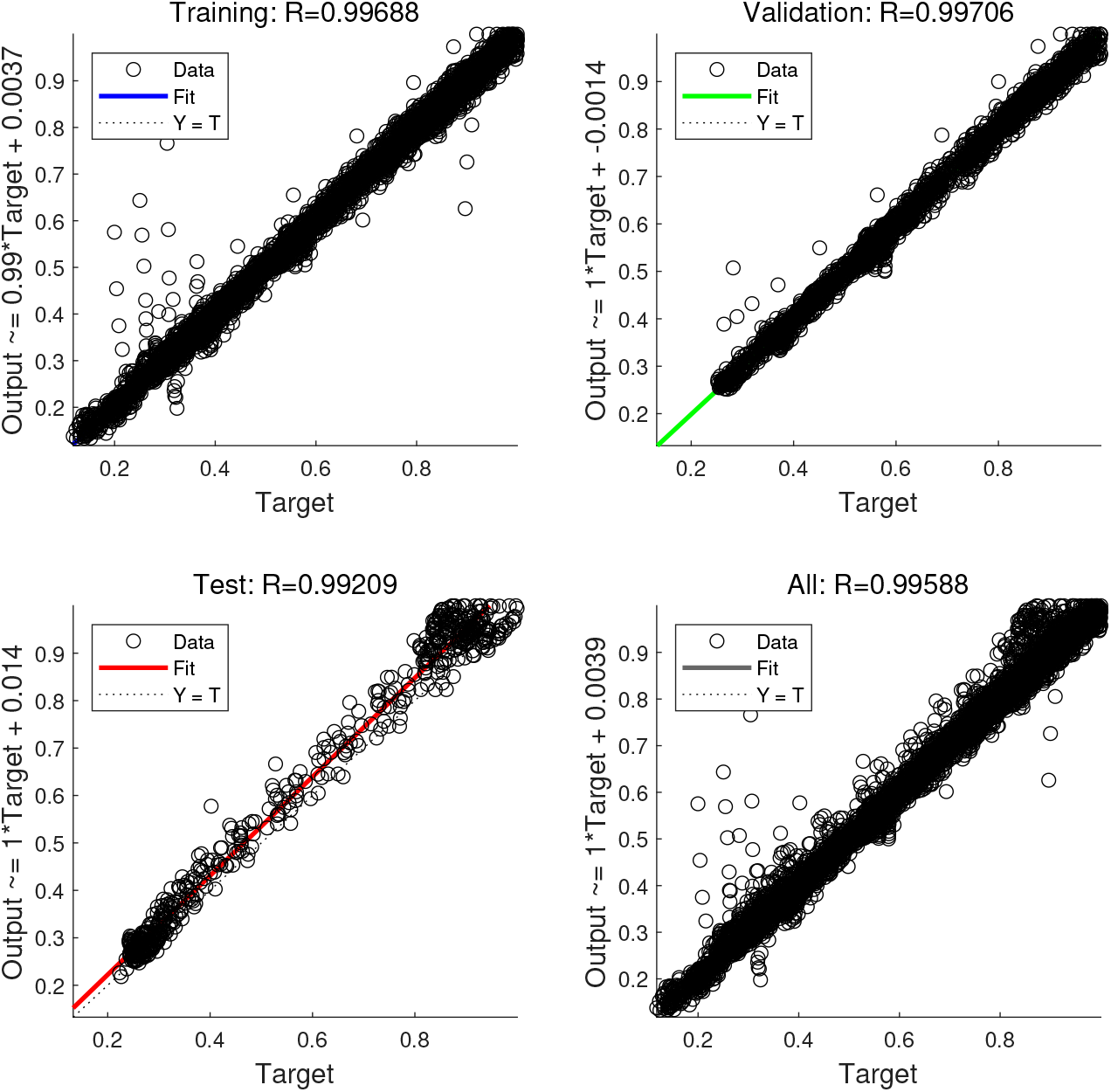
The machine learning training, validation, and testing regression of a general model of RNN MuscleNET with filtered sEMG and delayed kinematic signals input and joint angle output. The test subject did not participate in training and validation.

For stochastic and time-varying sEMG signal inputs in the machine learning model, it is essential to have rich data to extract and validate the model and configure the model architecture. Designing a machine learning model with many layers and millions of variables requires very large datasets. The combination of training, validation, and testing datasets with 1530000 pairs of inputs and outputs were sufficient for MuscleNET (a shallow neural network with three layers and a deep neural network with eight convolutional layers and a small filtering size).

### 4.2. Models Comparison

The comparison of models focuses on their configuration identity, input types, outputs, and the delay time. For consistent accuracy comparison, the outputs were normalized by the maximum value for each subject. The comparison of 80 configurations allows consistent comparison and directions for future research. Based on the comparison of results in Table 3, the preferred model for (I) offline simulation of a musculoskeletal model and (II) real-time control of a biomechatronic device is presented.

**Table 3.**
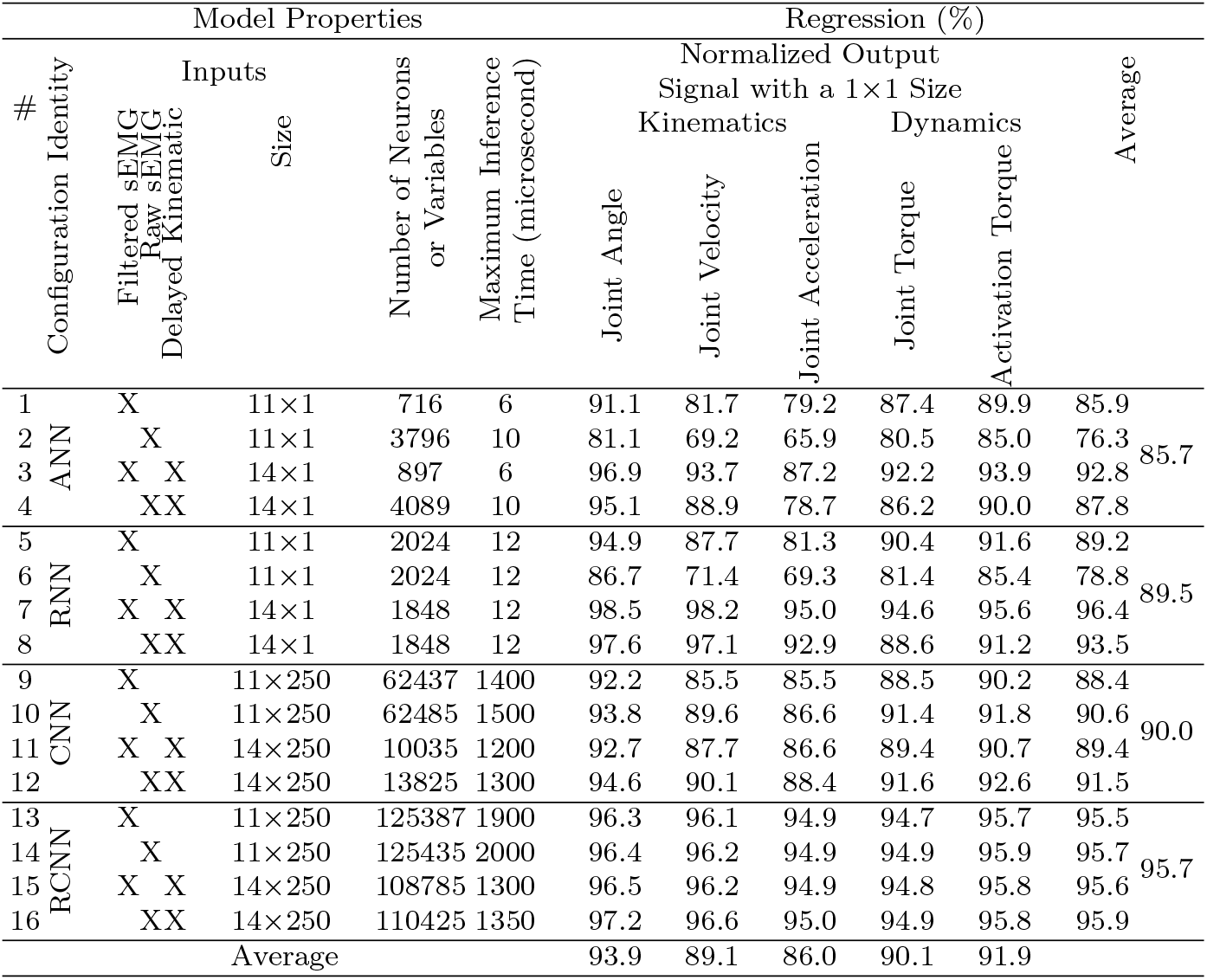
Comparison of regression accuracy for 80 models: 16 MuscleNET configuration times 5 output signals.

#### 4.2.1. Models’ Topology

From a topology viewpoint (Table 3), the machine learning method with the recurrent topology or the feedback outperforms the feed-forward mapping (RNN and RCNN had average regression accuracy of 89.5% and 95.7%, respectively, while ANN and CNN had lower average regression accuracy of 85.7% and 90.0%, respectively). Previously, Chen et al. [13] reported that the recurrent-based models, RCNN consisting of Long Short-Term Memory (LSTM) networks, were much better than a CNN consisting of a Fully Connected (FC) network, which agrees with our conclusion based on current results (Table 3). The reason behind this is associated with the relation of sEMG to the motion. According to the active and passive muscle tension behavior utilized for the Hill-type muscle dynamic model [4], the relationship is not straightforward and has a second-order differential equation. The recurrent topology of the RCNN uses the previous output signals as the inputs; therefore, it can model a higher-order differential equation.

#### 4.2.2. Models’ Outputs

In terms of the models’ outputs (Table 3), accuracy varied. In comparison to Bao et al. [33] which used a CNN for regression of only wrist joint angle, we have evaluated different output signals (Kinematic and Dynamic Biomechanical Variables) and calculated relative accuracy, which helps improve the future application of MuscleNET for myoelectric-based control. The general accuracy of the outputs in ascending order is: output joint angle, activation torque, joint torque, output joint velocity, and output joint acceleration (with average regression accuracy of 93.9%, 91.9%, 90.1%, 89.1%, and 86.0%, respectively). This order concurs with mathematical muscle models. The muscle pennation angle, muscle wrapping, muscle-tendon length, and passive muscle torque are functions of the joint angle [4]. In addition, the relation of muscle velocity and activation dynamic of muscle has been seen in the MTG (muscle torque generator) models or activation torque [48]. The maximum muscle force was relevant to the joint torque [4]. Thus, the first three dominant output signals correspond to the primary variables in classical muscle models. Thus, we propose using the sEMG signal to control biomechatronic devices using the joint angle or activation torque. In this regard, since most robotic servo motors have a PID position control, using the sEMG as the input of the control algorithm and the joint angle as a command to the low-level actuator control is more straightforward. Another method is using a force/torque-based control algorithm for biomechatronic devices. With the sEMG used as the input, the machine learning model can estimate the activation torque using an MTG model [50]. The joint torque can be calculated and commanded to the low-level control loop or the robot actuator. The activation torque’s superiority over joint torque arises from consideration of the passive and active torque impact of muscle in the MTG model [48, 50].

#### 4.2.3. Models’ Inputs

From the kinematic input standpoint, using the delayed kinematic signals along with the sEMG signals positively affected the model’s accuracy (using delayed kinematic signals resulted in an average regression accuracy of 92.9% while not using those signals resulted in a lower average regression accuracy of 87.6%). It is noteworthy to mention that the Hill-type muscle model’s inputs [4] are the joint angle, velocity, and acceleration, along with the sEMG signal. Thus, using the kinematic signals along with the sEMG signal, for the first time, is novel and increased model accuracy. Indeed, incorporating the delayed kinematic signals decreased the regression error from 50% to 20%. For example, using the delayed kinematic signals in row 12 in Table 3 provides more accuracy than row 10, which results from the model proposed by Ameri et al. [35]. Using the delayed signals instead of the current signals reflects the need to consider electrical and computational delays. Moreover, for real-time control of an actual bio-robot, the delay must be considered in the modeling. For the simulation of the musculoskeletal model, delayed signals are unnecessary.

From the sEMG input perspective, the result provided two distinct conclusions regarding other input conditions: raw or filtered sEMG signal. First, for the ANN and RNN models, the models’ accuracy was much better when the inputs were filtered signals (filtered sEMG signals yielded average regression accuracy of 91.1%, while raw signals yielded average regression accuracy of 84.4%). Hence, the ANN and RNN models should not be used for filtering the raw signals. The results revealed the capability of the convolutional models in filtering the signals and confirm the superiority of using raw sEMG signals as the inputs of the CNN and RCNN models.

#### 4.2.4. Models’ Delay

Since real-time control of biomechatronic devices requires a minimal delay, this factor should be considered. Comparison of the system’s delay is controversial since the signal filtering delay should be considered as well. Subsequently, filtering the raw sEMG signal has a specific delay due to low-pass filtering. The delay in filtering the raw signals should be considered with the model delay to compute the total delay. In this regard, the models’ delay with the filtered sEMG signals exceeds expectations. All the control methods had an operation delay of less than 2 milliseconds (Table 3) because of shallow configuration (not having too many layers), which is quick for real-time control purposes. Compared to sEMG frequency-based input [36, 37], the MuscleNET is much faster and works in real-time since prior research [36, 37] initially converted sEMG signal to sEMG frequency bands, which is an offline and time-consuming conversion. Moreover, sEMG-based classification of hand or wrist gestures [5, 6, 12, 17, 31, 32] requires additional sEMG history compared to RNN MuscleNET which mainly uses the signal value and a faster regression. The Ameri et al. [35] model also has five times more variables than CNN MuscleNET with delayed kinematic input, indicating that MuscleNET incurs less computation cost and has less delay than that model [35].

### 4.3. Testing and Demonstration

These findings motivated re-learning of two specific models and comparison of their results. In those models, the outputs were the joint angle and activation torque. The two models were (I) RNN with filtered sEMG signals plus delayed kinematics and (II) RCNN with raw sEMG signals plus delayed kinematics. For re-training of the RNN, the maximum elapsed time was set to 12 hours. For re-training of the RCNN MuscleNET, the maximum epoch was set to 1000. The rest of the training options were the same as the previous training step. As an example, the RCNN training and validation error rates for 1000 iterations are presented (Figure 14). The performance of RNN and RCNN MuscleNET for (a) joint angle and (b) activation torque trajectory with a user dataset that had not been used for training is shown in Figure 15. The RCNN MuscleNET followed the target values, which were the normalized joint angle and joint torque. Another random subject data was used to evaluate the model (Figure 16). This complete estimation of the biomechanical signals of random subjects, who did not participate in training and validation of the model, had 96.9% accuracy. The result showed that the novel model configuration, using delayed kinematic signals, the optimum model configuration, the given amount of data, and the optimum division for training, validation, and testing successfully achieved the goal. The amount of experimental data (1530000 pairs of inputs and outputs) was sufficient to reach this regression accuracy for estimation tests.

**Figure 14.**
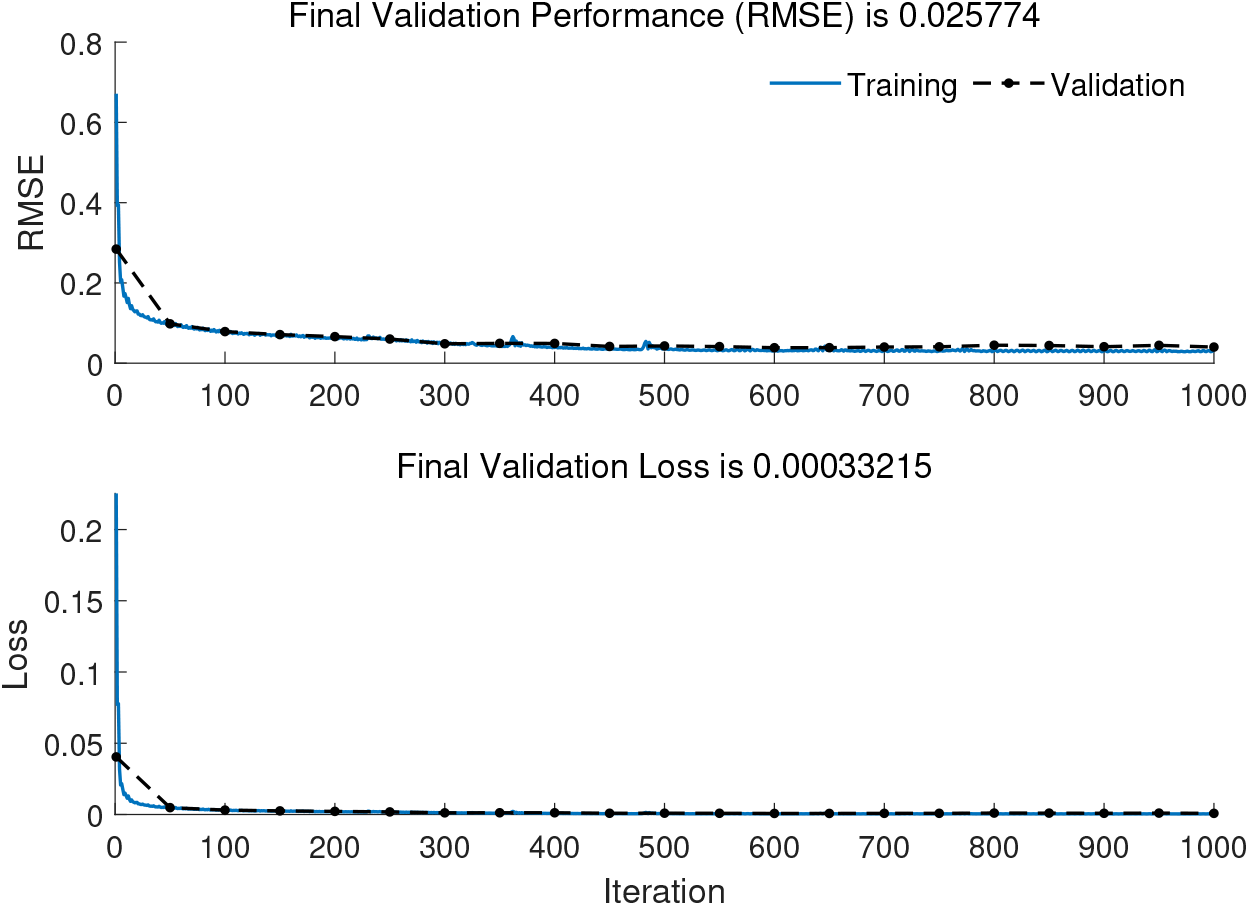
The RCNN training (solid) and validation (dashed) rates for 1000 iterations. However, the training procedure could be stopped at 500 iterations.

**Figure 15.**
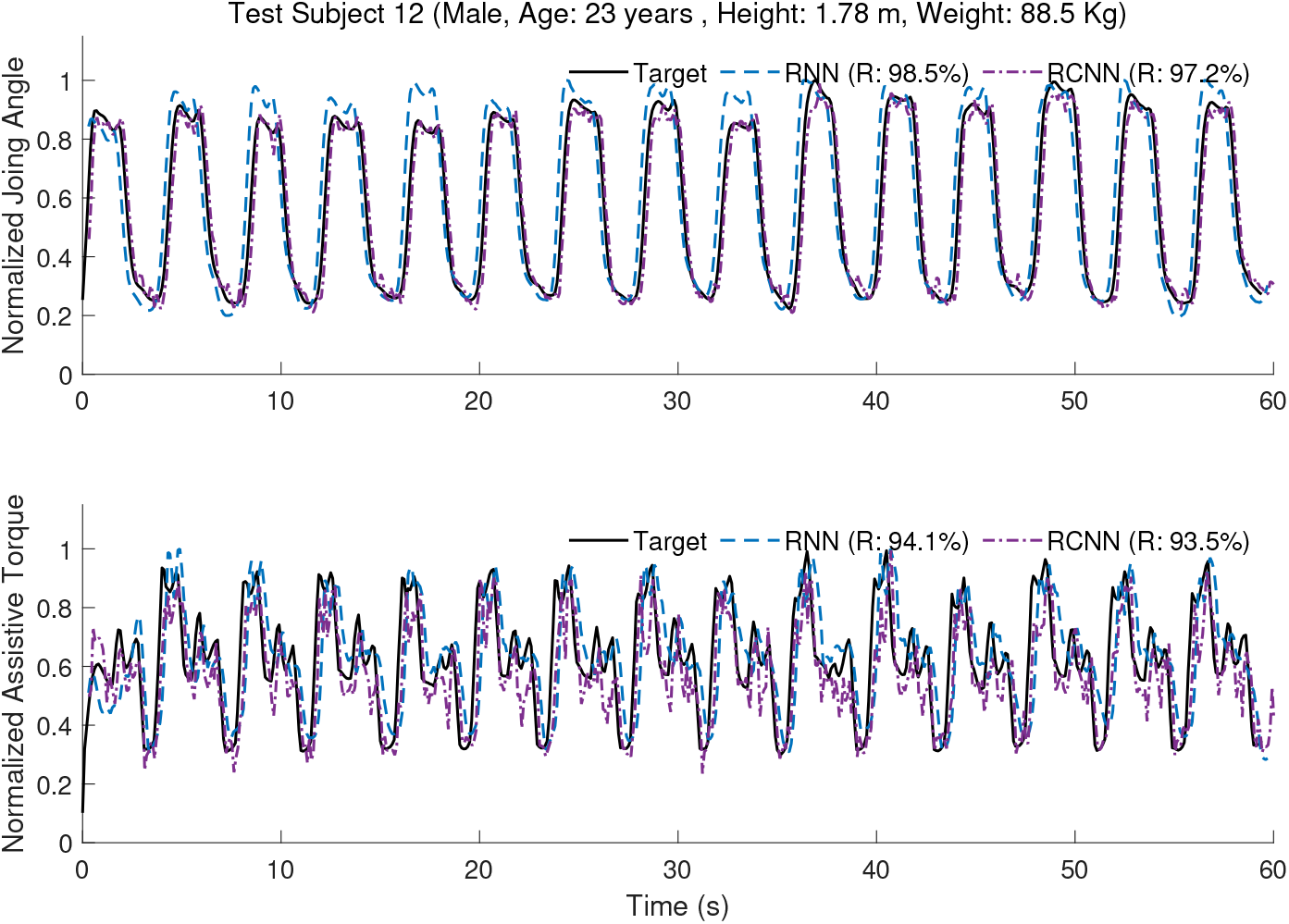
The performance of RNN and RCNN MuscleNET on random test data for users’ joint angle (top) and joint torque trajectory (bottom).

**Figure 16.**
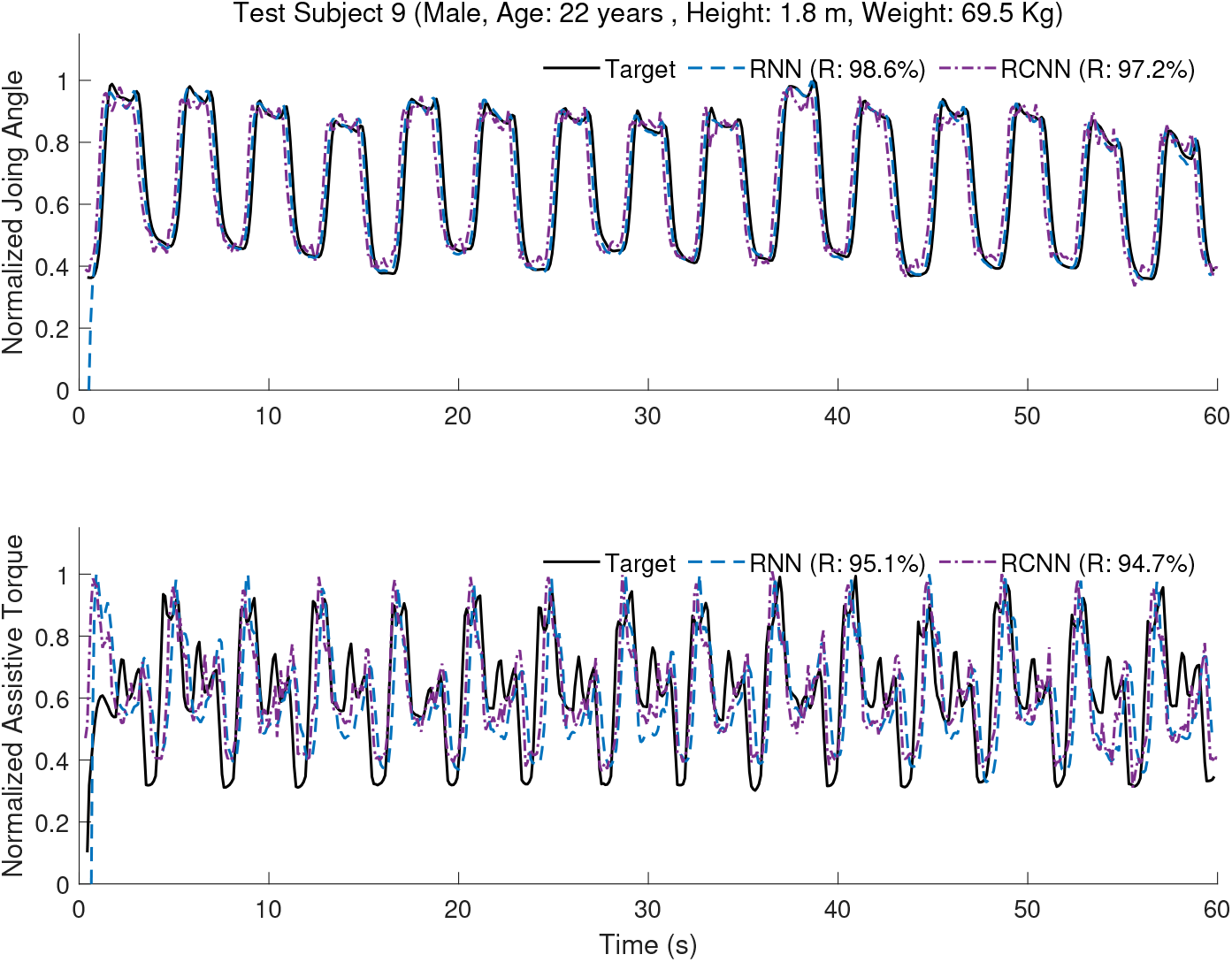
The test of RNN and RCNN MuscleNET for second random non-used user data for training users’ joint angle (top) and joint torque trajectory (bottom).

### 4.4. Application

Two specific applications for (i) RNN MuscleNET and (ii) RCNN MuscleNET emerge from these findings:

i. Using RNN MuscleNET with inputs of filtered sEMG and delayed kinematic signals could replace muscles in musculoskeletal models. The output of the MuscleNET can be joint elevation torque or activation torque, and the output of the MuscleNET should be used as the input of the skeletal dynamic model. MuscleNET and the skeletal model’s total system can be used in the forward dynamic simulation of a musculoskeletal model (for example, for clinical or sport engineering purposes) (Figure 17).
ii. For biomechatronic purposes, the RCNN MuscleNET, with raw sEMG and delayed kinematic signals, can be used as an interpreter of the sEMG signals. The output of RCNN MuscleNET is commanded to the low-level control loop of biomechatronic systems (Figure 18). Applications of the method may include controlling assistive/resistive robots, exoskeletons, or prostheses [1, 58].

**Figure 17.**
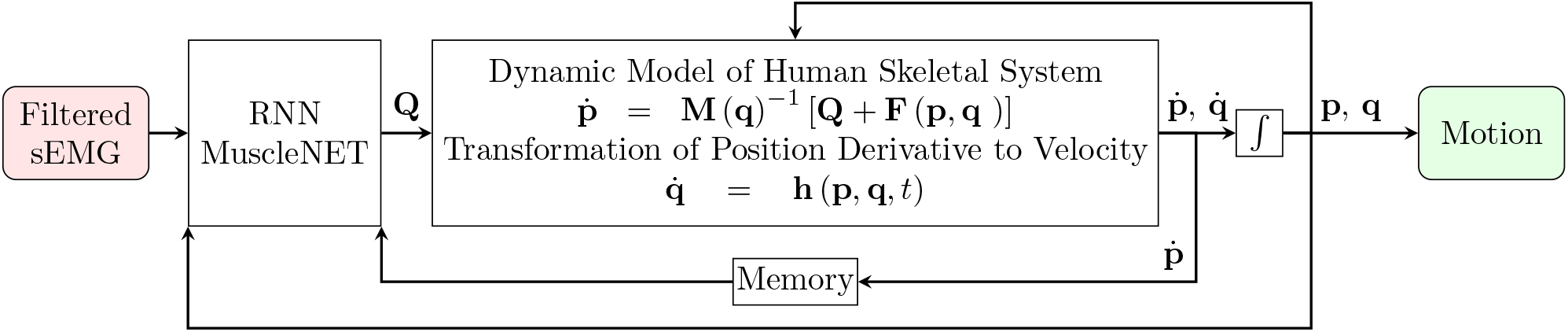
Schematic of a musculoskeletal model simulation (for example, for sport engineering purposes or musculoskeletal analysis). MuscleNET acts as a muscle model using the filtered sEMG, joint position, velocity, and acceleration. MuscleNET’s output is the joint torque *τ_h_* supplied to the dynamic model for forward dynamic simulation.

**Figure 18.**
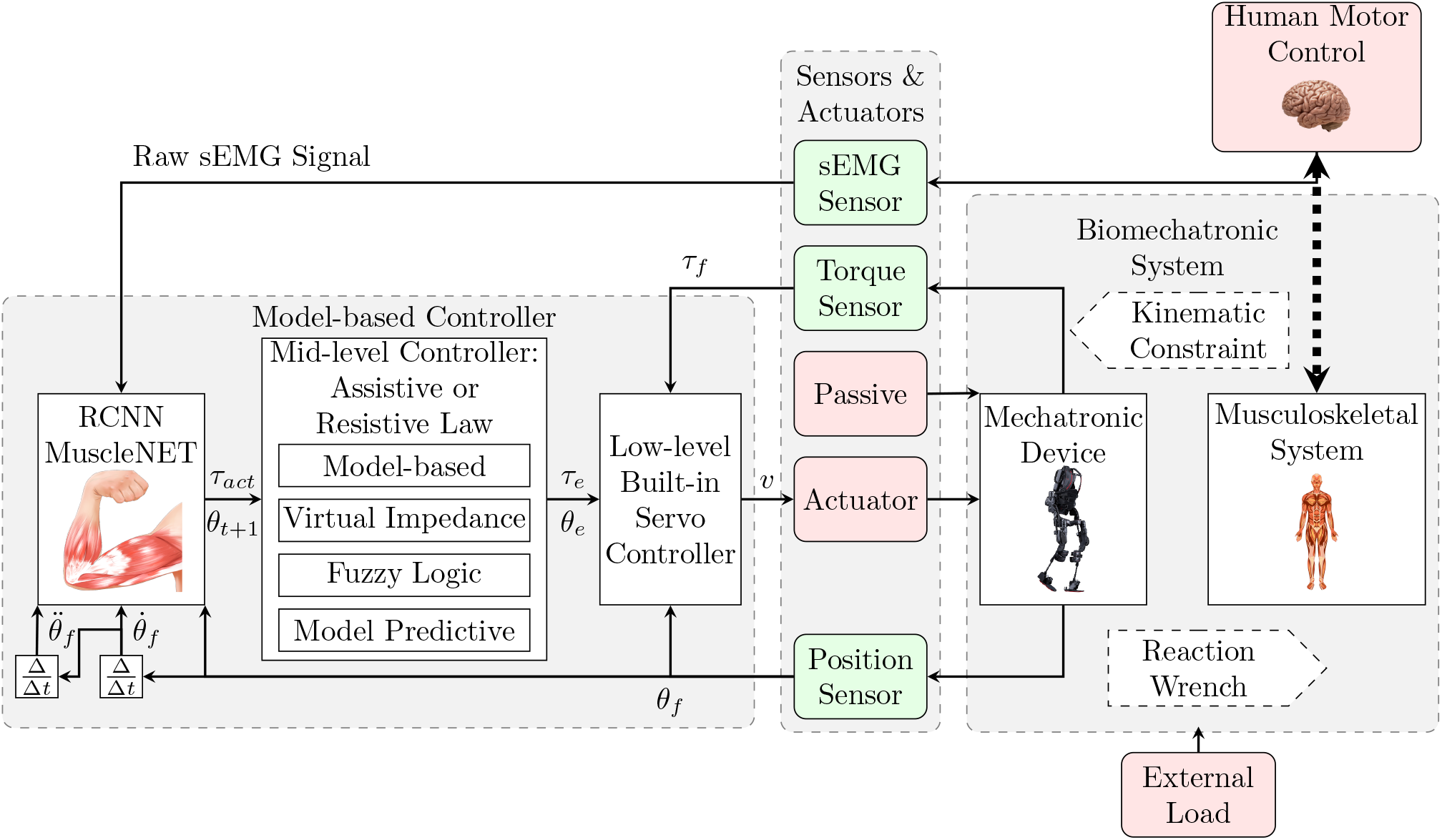
Schematic for controlling an exoskeleton, a prosthesis, or a rehabilitation robot using MuscleNET as a muscle model for converting the raw sEMG, joint feedback position *θ_f_*, velocity 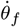, and acceleration 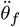 to activation torque *τ_act_*. The low-level controller is responsible for controlling the mechatronic system by desired torque *τ_e_* or angle *θ_e_* according to the output from mid-level controller.

## 5. Conclusion

An alternative solution to mathematical muscle modeling using regression RNN and CNN-based sEMG signals is proposed. The model’s performance and quality metrics were compared across 80 different machine learning regression-based schemes with various conditions. The conditions consisted of (I) different kinds of outputs: 3 kinematic signals (joint angle, velocity, and acceleration) and 2 dynamic signals (joint torque and activation torque), (II) different input conditions: using delayed kinematic signals, raw, or filtered sEMG signals, and (III) different kinds of neural network configurations: ANN, RNN, CNN, and RCNN. The two models of (A) RNN with delayed kinematic signal and filtered sEMG signals inputs and (B) RCNN with delayed kinematic and raw sEMG signal inputs outperformed the other schemes in terms of estimation accuracies. The high performance of the recurrent-based models demonstrated their ability to learn necessary motor information from sEMG signals. Further, the signal filtering capability of the CNN-based models was established. MuscleNET was more accurate in estimating joint angles and activation torque signals than joint velocity, joint acceleration, and joint torque.

Future work should study the robustness of MuscleNET to factors such as electrode shift and electrodes that disconnect during a motion. Moreover, the practical application of MuscleNET to control an active exoskeleton or prostheses is currently being pursued by the authors, and should lead to refinements in the method.

## Acknowledgments

The authors acknowledge funding of this research by the Canada Research Chairs program.

